# CTCF blocks anti-sense transcription initiation at divergent gene promoters

**DOI:** 10.1101/2021.10.30.465508

**Authors:** Jing Luan, Camille M. Syrett, Marit W. Vermunt, Allison Coté, Jacob M. Tome, Haoyue Zhang, Anran Huang, Jennifer M. Luppino, Cheryl A. Keller, Belinda M. Giardine, Shiping Zhang, Margaret C. Dunagin, Zhe Zhang, Eric F. Joyce, John T. Lis, Arjun Raj, Ross C. Hardison, Gerd A. Blobel

## Abstract

Transcription at most promoters is divergent, initiating at closely spaced oppositely oriented core promoters to produce sense transcripts along with often unstable upstream antisense (uasTrx). How antisense transcription is regulated and to what extent it is coordinated with sense transcription is largely unknown. Here by combining acute degradation of the multi-functional transcription factor CTCF and nascent transcription measurements, we find that CTCF specifically suppresses antisense but not sense transcription at hundreds of divergent promoters, the great majority of which bear proximal CTCF binding sites. Genome editing, chromatin conformation studies, and high-resolution transcript mapping revealed that precisely positioned CTCF directly suppresses the initiation of uasTrx, in a manner independent of its chromatin architectural function. Primary transcript RNA FISH revealed co-bursting of sense and anti-sense transcripts is disfavored, suggesting CTCF-regulated competition for transcription initiation. In sum, CTCF shapes the transcriptional landscape in part by suppressing upstream antisense transcription.

## Main Text

Divergent transcription at active promoters is prevalent among eukaryotes, producing upstream antisense transcripts (uasTrx) that are rapidly processed and tend to be short lived^1-3^. Divergent promoters are nucleosome-depleted region densely occupied by transcription factors. They typically harbor two distinct core promoters positioned in inverted orientations, instructing the assembly of separate transcription pre-initiation complexes (PICs) that transcribe along opposite DNA strands^4-7^. Transcriptional outputs by divergent promoters in both orientations are generally concordant, suggesting co-regulation^1,2,8-10^. In certain cases, however, sense and antisense transcription appears to be anti-correlated^11^. It thus remains unclear whether and how divergent transcription is coordinated spatially and temporally. On one hand, divergent transcription may be cooperative, as simultaneous presence of two PICs may help maintain nucleosome-depleted regions and allow for efficient transcription factor recruitment^5,12^. On the other, divergent PICs may compete for common transcription activators or physical space, thus rendering co-occurrence unfavorable^13^.

CTCF (CCCTC-binding factor) was first identified as a transcription factor and was later recognized to also shape genome topology together with the cohesin protein complex^14^. CTCF depletion is known to cause genome-wide architectural perturbation but limited changes in the transcription of coding genes^15-23^. However, the mammalian genome is ubiquitously expressed, producing abundant noncoding transcripts that have now gained increasing recognition as functional^24^. Whether and how CTCF affects the noncoding transcriptome remains to be explored experimentally.

We performed precision nuclear run-on sequencing (PRO-seq) in the mouse murine erythroid cell line G1E-ER4 in which both *CTCF* alleles have been modified to bear an auxin-inducible degron (AID) that allows for rapid CTCF degradation^23^. PRO-seq interrogates nascent transcription in a strand-specific manner at single base-pair resolution^25^. Overall, we observed limited perturbation of annotated transcripts after acute CTCF depletion^23^. Notably, however, at 376 active promoters we observed a significant increase in uasTrx production (Fig. 1a-c and Supplementary Data 1). These changes were corroborated by ChIP-seq (chromatin immunoprecipitation sequencing) of RNA polymerase II subunit A (POLR2A) and RT-qPCR at 3 select loci (Extended Data Fig. 1a,b). UasTrx were heterogenous in size, with the median being 1956 nucleotides (Extended Data Fig. 1c). The most 5’ ends of these transcripts initiated upstream of sense transcription start sites (TSSs). The average distance of the most frequently used uasTrx start sites from sense TSSs was approximately 100 bp (Fig. 1d), which is similar to that of divergent promoters found in other mammalian cells where this distance was ∼110 bp^5,13^. CTCF depletion led to increases only in the antisense direction leaving sense transcription ostensibly unperturbed, suggesting that CTCF promotes the directionality of divergent promoters by exerting strand-specific transcription repression (Fig. 1d and Extended Data Fig. 1b,d,e,f).

**Fig. 1.**
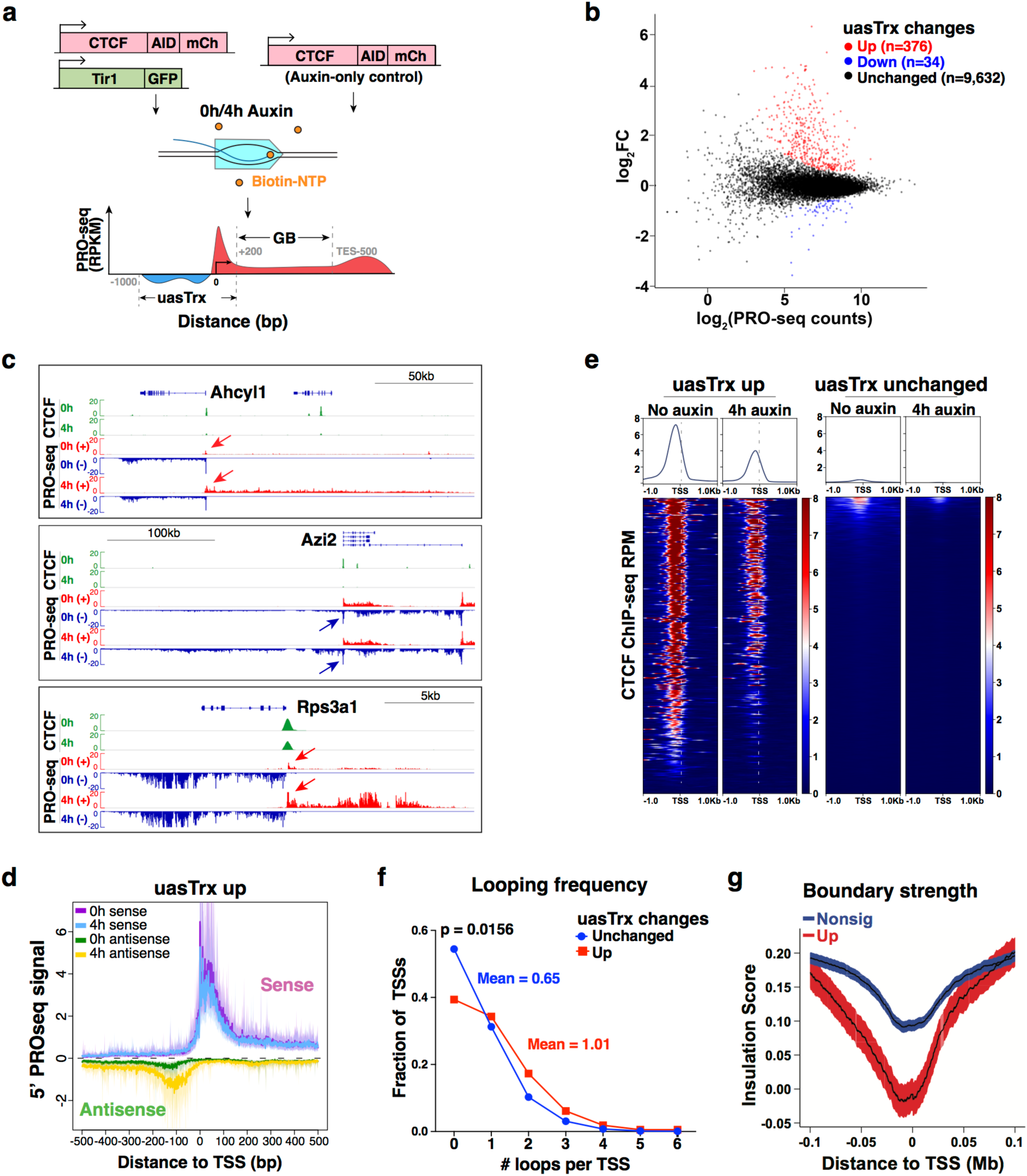
Transient CTCF depletion leads to widespread antisense transcription up-regulation at divergent promoters. **a**, Schematics of PRO-seq experiment and quantification strategy. **b**, PRO-seq MA plot of control versus CTCF-depleted cells on the antisense strand (−1000bp to +200 relative to annotated TSS) in G1E-ER4s. Differentially expressed transcripts highlighted in color. **c**, Genome browser views of CTCF ChIP-seq and PRO-seq signals at *Ahcyl1, Azi2* and *Rps3a1* loci. Arrows point to uasTrx, with colors indicating strandedness. **d**, Metaplot of sense and antisense 5’ PRO-seq signals at activated uasTrx, centered at annotated TSSs and plotted with respect to sense orientation. Solid lines and shades show bootstrapped estimates of average signals and the 12.5/87.5 percentiles, respectively. **e**, Row-linked heatmaps showing CTCF occupancy at active promoters (up n=376; unchanged n=9,632), grouped by uasTrx changes upon CTCF depletion, sorted by occupancy level and shown with respect to sense orientation. **f**, Frequency of looping interactions engaged by all gained and unchanged uasTrx. *P* value calculated by Wilcoxon signed-rank test. **g**, Averaged insulation score centered at annotated TSSs over 0.2Mb window, grouped by uasTrx changes, and plotted with respect to sense orientation.

Different terms have been used to describe antisense transcription from divergent promoters in eukaryotes, including cryptic unannotated transcripts (CUTs), stable unannotated transcripts (SUTs), and Xrn1-sensitive unstable transcripts (XUTs) in yeast, as well as PROMoter uPstream Transcripts (PROMPTs) and “upstream divergent transcripts” in higher eukaryotes^1,3,9,26,27^. The uasTrx that we found to be repressed by CTCF may represent a subset of PROMPs/upstream divergent transcripts.

Promoters with up-regulated uasTrx are enriched with proximal (mostly <100bp from annotated TSSs) CTCF binding, reminiscent of an earlier finding observed across multiple human cell lines (Fig. 1e and Extended Data Fig. 2a)^28^. Notably, only a fraction of CTCF-bound promoters (319 out of 1,846) increased uasTrx production in response to CTCF loss, but those tended to have a stronger CTCF binding intensity (Extended Data Fig. 2b). However, CTCF binding reduction and uasTrx gains were only weakly correlated (Extended Data Fig. 2c). Because strong CBSs tend to be conserved across cell types^15,23^, we assessed CTCF occupancy across mouse tissues^29^. Indeed, CBSs at uasTrx regulatory sites were more tissue-invariant, indicating that uasTrx repression may be a conserved feature (Extended Data Fig. 2d). To assess whether CTCF functions in a similar way in other species and tissues, we measured uasTrx changes upon CTCF depletion in the human colorectal carcinoma cell line HCT-116 and again found 199 uasTrx to be up-regulated without significantly affecting sense transcription (Extended Data Fig. 3a). We also examined published data sets in mouse embryonic stem cells (mESCs) and observed a similar number of up-regulated uasTrx (Extended Data Fig. 4a,b)^30^. Antisense changes in both cell types were similarly associated with strong promoter-proximal CTCF binding (Extended Data Fig. 3b and Extended Data Fig. 4c) and a lack of sense perturbation (Extended Data Fig. 3c-f and Extended Data Fig. 4d-f). Lastly, up-regulated uasTrx in mESCs were silenced upon CTCF recovery following auxin removal (Extended Data Fig. 4b,g). Hence, CTCF represses uasTrx at numerous genes across multiple species and cell lineages.

Because promoter-proximal CTCF only suppresses a subset of the uasTrx, we examined features that determine uasTrx regulation by CTCF. In addition to being enriched with strong CBSs, promoters with up-regulated uasTrx harbored high levels of cohesin, a protein complex central to genome folding^31,32^, compared to those that were unchanged upon CTCF depletion (Extended Data Fig. 5a). In addition, they are enriched at chromatin loop anchors and chromatin domain boundaries (Fig. 1f,g and Extended Data Fig. 5b,c). The associated sense transcripts also tend to be housekeeping genes, which are frequently found at domain boundaries^33^ (Extended Data Fig. 5d). Finally, in yeast, chromatin looping (“gene loops”) was implicated in the control of transcription directionality^34^. Therefore, we interrogated the possibility of CTCF controls transcription directionality by regulation architectural functions.

We first determined whether the repressive effects of CTCF on uasTrx were direct by disrupting CTCF binding at TSS-proximal CTCF binding site (CBS) at three loci (Fig. 2a), *Ahcyl1, Azi2* and *Rps3a1*, through CRISPR/Cas9-mediated genome editing (Fig. 2b, Extended Data Fig. 7a and Extended Data Fig. 7-8a)^35^. Upon disruption of TSS-proximal CTCF binding, uasTrx production became elevated while sense transcription remained unperturbed, demonstrating that CTCF binding directly constrains uasTrx production (Fig. 2c-e, Extended Data Fig. 6a,b, Extended Data Fig. 7a-d,f and Extended Data Fig. 8b-d).

**Fig. 2.**
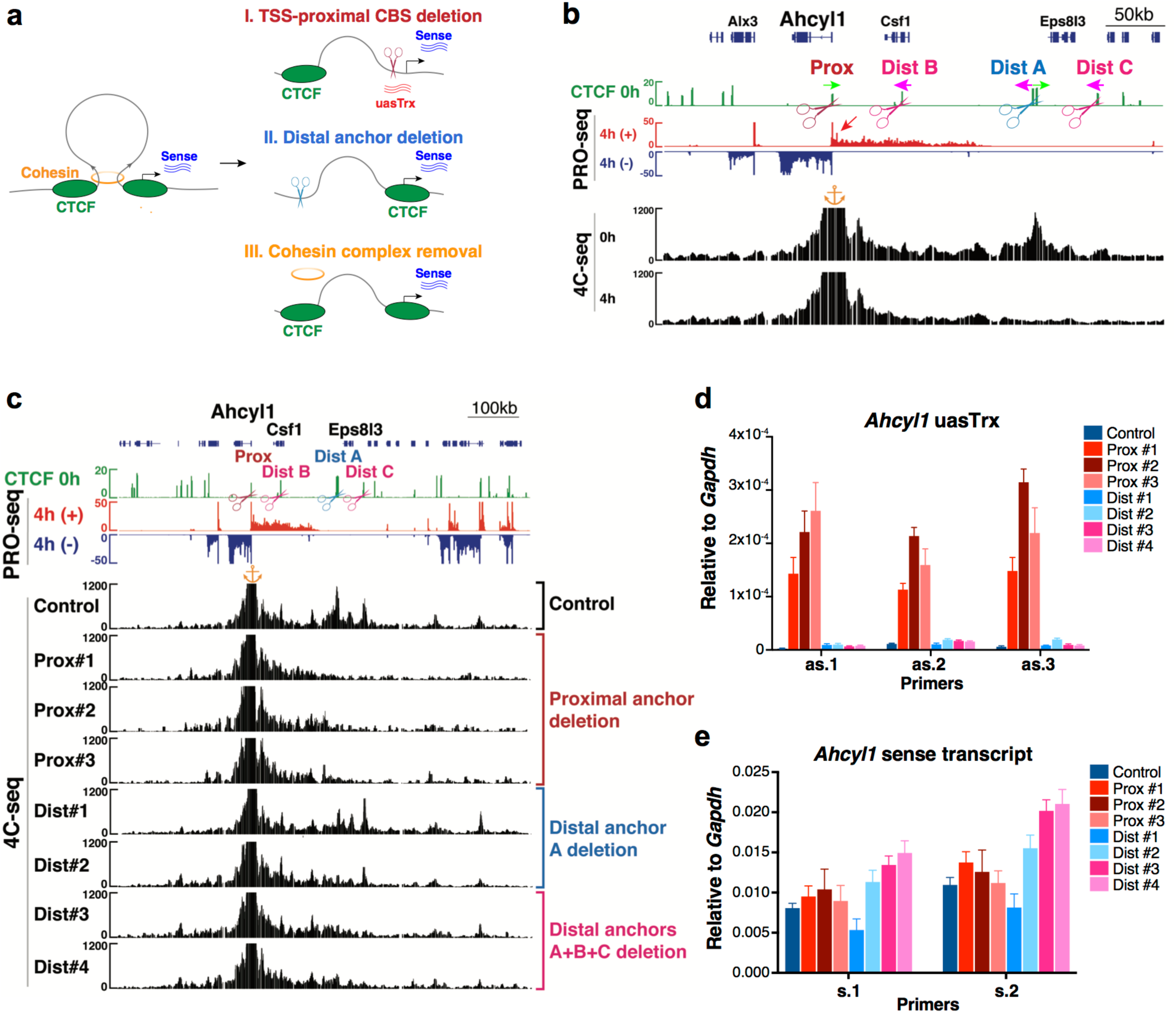
CTCF inhibits uasTrx directly and proximally, independent of its architectural functions. **a**, Illustration of experimental strategy and summarized findings in **Fig. 2** and **Extended Data Figs. 6-9. b**, Genome browser views of CTCF ChIP-seq, PRO-seq and 4C-seq signals at *Ahcyl1*. 4C-seq anchored at *Ahcyl1* promoter. Colored arrows above ChIP-seq track indicate CTCF motif directionality. Red arrow points to elevated uasTrx after CTCF depletion. Scissors point to regions disrupted by CRISPR/Cas9-mediated genome editing, one at CBS proximal to *Ahcyl1* promoter and the others at a distal CBSs engaging in loop contacts with the promoter. Orange anchor indicates 4C-seq viewpoint. **c**, Genome browser tracks of bulk CTCF ChIP-seq and PRO-seq and representative 4C-seq profiles of control and edited clones with indicated regions disrupted. Similar observations were made in 2 additional independent 4C-seq experiments and not shown. Orange anchor indicates 4C-seq viewpoint. **d**, RT-qPCR of *Ahcyl1* uasTrx in control and edited clones. Transcripts were normalized to *Gapdh* (error bar: SEM; n=3-4). **e**, Same as (**d**) but of nascent *Ahcyl1* sense transcripts. Prox, TSS-proximal CBS. Dist, distal anchor.

Most chromatin boundaries are occupied by CTCF; however, a large fraction of CTCF sites is not associated with domain boundaries or measurable chromatin loops^33^. We thus employed 4C-seq (Circularized Chromosome Conformation Capture sequencing) to determine whether CTCF-bound promoters engage in long-range looped interactions^36^. We focused on the 2 loci, *Ahcyl* and *Azi2*, where uasTrx was strongly and directly suppressed by CTCF, and found significant looping interactions with distant CBSs (Fig. 2b and Extended Data Fig. 7a). Upon CTCF depletion, these loops were strongly diminished, indicating that CBSs are indeed involved in architectural functions at these 2 genes. In light of prior studies in yeast invoking gene looping as a mechanism to constrain uasTrx, we assessed whether CTCF’s architectural function constrains uasTrx production^34^. Inspection of the 4C-seq tracks identified the most prominent loop anchors, which we disrupted via CRISPR-Cas9 mediated genome editing in a manner that preserved promoter-proximal CTCF binding. At the *Ahcyl1* gene, deletion of the distal CTCF site (Dist A) that is associated with the most prominent loop promoter loop (Dist A) led to loss of 4C-seq contacts (Fig. 2c and Extended Data Fig. 6c-e) but no change in uasTrx production (Fig. 2d,e). However, since some contacts remained, we removed two additional CBSs at 4C-seq contact sites (Dist B and Dist C), which further reduced interactions with the promoter proximal CBS (Fig. 2c). None of these perturbations increased uasTrx production (Fig. 2 d,e). At the *Azi2* locus, deletion of distal loop anchors (Dist A and Dist B) but not promoter-proximal CBS led to significant architectural perturbations (Extended Data Fig. 7b-e). In contrast to promoter-primal CBS removal, Dist A/Dist B deletions failed to increase uasTrx production (Extended Data Fig. 7f). Of note, neither CTCF depletion nor CBS removal at the promoters of the *Ahcyl1* and *Azi2* genes detectably increased contacts between uasTrx promoters and surrounding putative enhancers (not shown). This argues against promoter-proximal CBSs functioning as enhancer blocking insulators. Together, these results separate the uasTrx repressive effects of CBSs from their architectural involvement at these loci.

Promoter-proximal CTCF sites involved in inhibition of uasTrx generation are enriched for cohesin (Extended Data Fig. 5a). As an independent means to assess a possible role of CTCF/cohesin-associated loops in regulating uasTrx production, we globally disrupted looped contacts by depleting Nipbl in HCT-116 cells, a cohesin-loading factor^37^, and interrogated transcriptional changes. PRO-seq experiments in Nipbl deficient cells revealed minimal uasTrx upregulation (Extended Data Fig. 9a). Finally, we analyzed published data sets in HCT-116 cells in which transient depletion of the cohesin component Rad21 was previously shown to cause genome-wide chromatin organization disruption^38^. Again, we did not observe strand-specific uasTrx changes. Instead, hundreds of genes underwent concomitant changes in both sense and antisense directions (Extended Data Fig. 9b-e), and were not enriched with strong CTCF or Rad21 binding at their promoters (Extended Data Fig. 9f). Together, three orthogonal approaches demonstrate that CTCF inhibits uasTrx directly and proximally, and independently of its architectural functions.

The process of transcription involves multiple steps, including initiation, pausing of RNA polymerase II (Pol II) after transcribing the first 20-60 nucleotides, and release of Pol II into the gene body (GB). CTCF was previously reported as capable of repressing pause-release in the sense direction^39^ and was also implicated in impeding Pol II elongation in the GB^40,41^. To determine the CTCF-controlled step(s) in uasTrx transcription, we took advantage of the single base-pair resolution afforded by PRO-seq and examined the distribution of CTCF motifs relative to the 5’ and 3’ PRO-seq signals which allows assessment of transcription initiation and stalling, respectively. Only active promoters with proximal (±100bp relative to TSS) CTCF binding sites harboring high-confidence CTCF motifs (motif score>75) were included in the analysis to ensure precise prediction of CTCF positioning (Extended Data Fig. 10a and Supplementary Data 2). Changes in transcription initiation and stalling would be expected to give rise to distinct PRO-seq patterns. Specifically, blockade of Pol II processivity would show significant accumulation of 3’ PRO-seq signals (i.e. paused Pol II) upstream of CTCF motifs, which would then get released upon CTCF depletion (Fig. 3a, “stalling”). Release from CTCF-mediated blockade on transcription initiation, on the other hand, would reveal enrichment of 5’ PRO-seq signal extending from the motif to the end of uasTrx after CTCF removal (Fig. 3a, “Initiation blockade”).

**Fig. 3.**
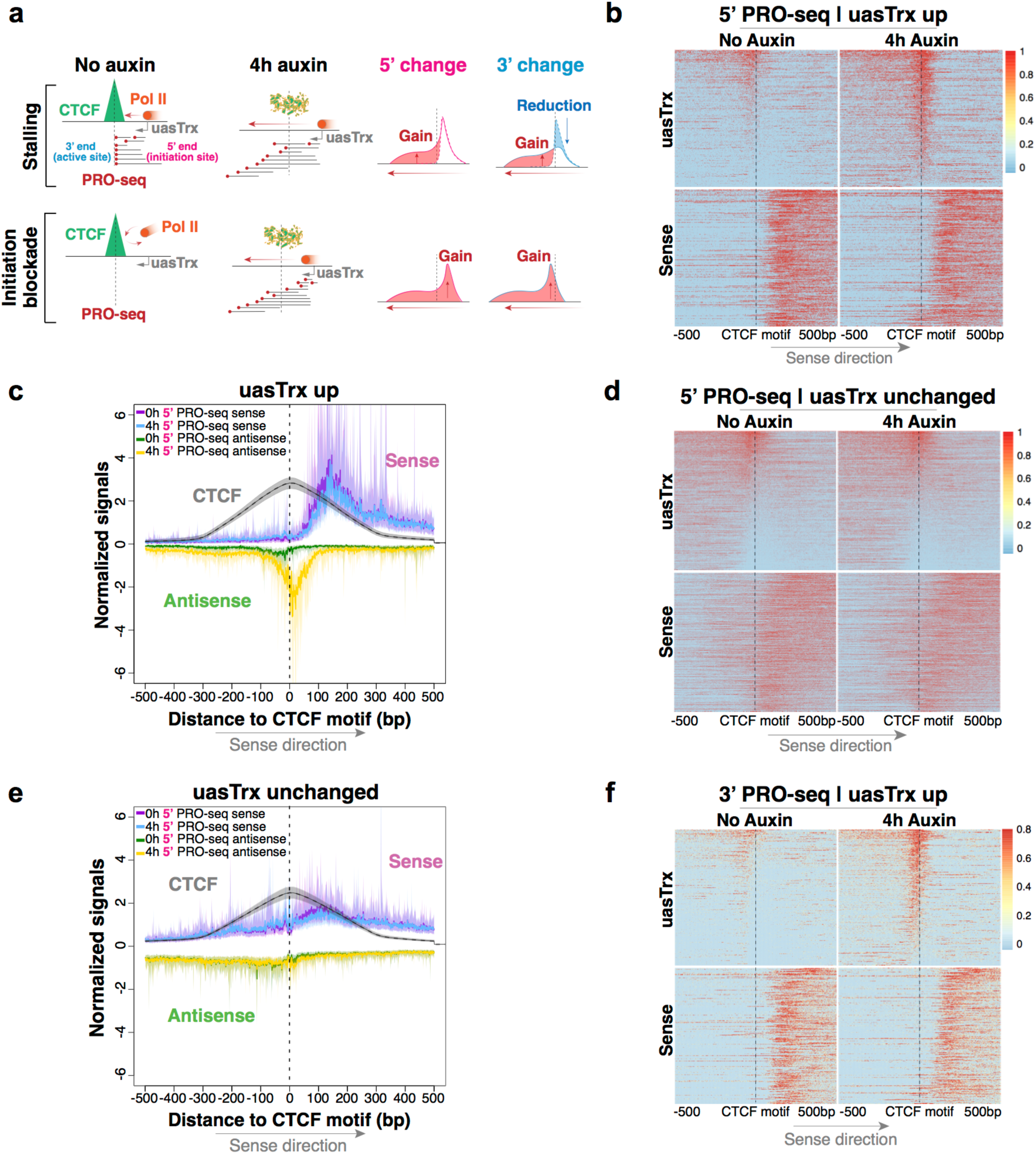
CTCF inhibits antisense transcription initiation through proximal binding. **a**, Model illustrating expected 5’ and 3’ PRO-seq distribution when CTCF blocks transcription processivity or initiation. **b**, 5’ PRO-seq heatmap at affected active promoters (n=298) that exhibit proximal CTCF binding and high-confidence CTCF motif(s) (motif prediction score>75), centered at CTCF motifs, sorted by mean antisense signal densities over the center 200bp and shown with respect to sense orientation. Black line highlights CTCF motif locations. **c**, Metaplot centered by CTCF motifs summarizing 5’ PRO-seq and CTCF signals shown in **b**. Solid lines and shades show bootstrapped estimates of average signals and the 12.5/87.5 percentiles, respectively. **d**, Same as (**b**) but at unaffected promoters (n=1,201) that satisfy the same CTCF criteria. **e**, Same as (**c**) but summarizing sites in **d. f**, Same as (**b**), but plotting 3’ PRO-seq signals at activated uasTrx.

The measured 5’ PRO-seq changes triggered by CTCF loss indicate that CTCF impacts antisense transcription initiation (Fig. 3b). Strikingly, CTCF is consistently positioned ∼20bp downstream of uasTrx initiation sites at affected promoters, reminiscent of a previous observation that CTCF tends to reside at the borders of transcription initiation clusters^42^ (Fig. 3b,c). This distinct spatial arrangement is in stark contrast to the much more variable distribution around unperturbed promoters (Fig. 3d,e). A fraction (120 of 1201) of the unperturbed promoters did have CBSs downstream proximally (Extended Data Fig. 10b, “Downstream proximal”). However, a closer look revealed an upward trend of uasTrx production even though they were not included in the perturbed group because of thresholding (Extended Data Fig. 10c,d). Thus, uasTrx appears to be linked to a particular positioning pattern of CTCF. Finally, 3’ PRO-seq reads accumulated downstream, rather than upstream, of CTCF motifs, indicating that Pol II can successfully pass through CTCF without stalling (Fig. 3f and Extended Data Fig. 10e-g). Altogether, the evidence points to CTCF repressing uasTrx transcription through initiation inhibition rather than Pol II stalling, which is consistent with our recent observation that CTCF binding does not strongly interfere with Pol II processivity in the gene body^23^.

Transcription is known to occur in bursts, with burst frequency and amplitude subject to modulation^43-45^. To investigate the effects of CTCF on bursting, and whether sense and antisense transcription are coordinated, we employed single-molecule fluorescence in situ hybridization (smFISH) to quantify at the *Ahcyl1* and *Rps3a1* loci 1) transcription burst size (i.e. amplitude), 2) burst fraction (related to burst frequency), and 3) co-burst frequency (Fig. 4a). CTCF depletion led to no significant changes in burst fraction or size on the sense strand, consistent with bulk PRO-seq readouts. Antisense transcription, on the other hand, underwent significant increases in burst fraction but minimal changes in burst size, suggesting that CTCF mainly affects antisense burst frequency without altering sense transcription dynamics (Fig. 4b,c).

**Fig. 4.**
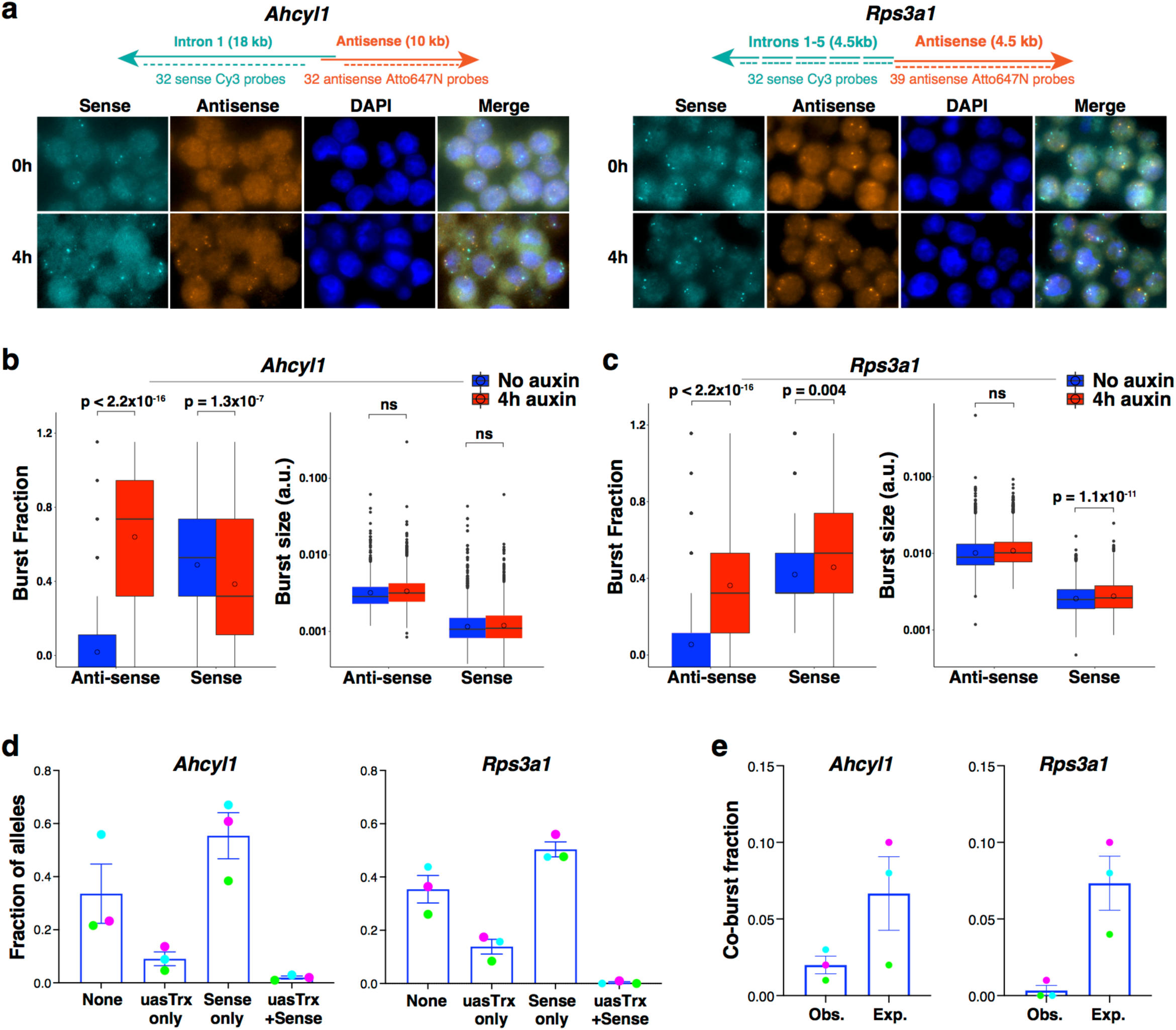
CTCF mainly regulates antisense burst fraction; sense and antisense bursts appear to compete temporally at divergent promoters. **a**, Top, maps of FISH probes targeting sense and antisense nascent transcripts at *Rps3a1* and *Ahcyl1* loci. Bottom, representative FISH images before and after CTCF depletion. **b**, Left, box plots showing antisense and sense burst fractions before and after CTCF depletion at *Ahcyl1*. Right, box plot showing antisense and sense burst sizes before and after CTCF depletion. n=3 biological replicates. *P* values were calculated by two-sample *t*-test. **c**, Same as (**b**) but at *Rps3a1*. **d**, Left, fraction of *Ahcyl1* alleles with 4 different sense/antisense bursting status at baseline (error bar: SEM; n=3). Right, same as left but at *Rps3a1*. Biological replicates matched by dot colors. **e**, Left, predicted and observed co-burst fraction at *Ahcyl1* at baseline (error bar: SEM; n=3). Right, same as left but at *Rps3a1*. Biological replicates matched by dot colors.

To interrogate sense/antisense burst coordination, we quantified the frequency at which both strands burst alone or together before and after CTCF depletion. At baseline, sense/antisense co-bursting occurred at a minimal number of alleles that is significantly less than expected (i.e. the product of sense and antisense burst fractions), suggesting that co-bursting is highly disfavored (Fig. 4d,e and Extended Data Fig. 11a). Upon CTCF removal, co-burst frequency increased significantly (Extended Data Fig. 11b) but still less frequently than would be expected if these events were independent of each other (Extended Data Fig. 11h). It is important to note that the results are confounded by the unexpectedly long half-lives (>4hr) of uasTrx at both loci (Extended Data Fig. 11c-g), which causes uasTrx transcripts to persist after completion of a burst, thus reducing temporal resolution of smFISH and inflating signal overlap. Regardless, sense and antisense bursts appear to be anti-coordinated temporally when transcribing from the same divergent promoter, suggesting competition between sense and antisense transcription initiation.

A variety of factors have been shown to affect uasTrx transcription, including R-loop formation, oncoprotein MYC, transcription elongation factor SPT6, transcription factor Rap1, looped contacts, histone modifications, and chromatin remodeling proteins (ex. Mot1, Ino80, NC2)^34,46-51^. In many instances, perturbations were accompanied by changes in the sense counterparts, which is in contrast to the present findings and suggests that CTCF functions through mechanism(s) distinct from those previously reported. On the other hand, the CAF-1 complex and histone H3K56 acetylation have been shown to suppress antisense transcription without significantly perturbing sense transcription in yeast^8^, but it remains to be tested whether a similar process is operational in mammalian cells and whether CTCF is involved.

Our smFISH results show that CTCF removal increases antisense burst fraction. Since CTCF can block enhancer-promoter contacts^52,53^ and since enhancers can increase burst fraction^54^, it is conceivable that CTCF loss leads to illegitimate enhancer contacts. However, we did not observe increased long-range contacts upon CTCF loss. Combined with our 5’ transcript mapping, this indicates that CTCF inhibits uasTrx production locally at the step of transcription initiation, possibly by preventing PIC formation. The dynamic relationship between sense and antisense transcriptional bursts has not been investigated previously. Single-molecule RNA-FISH at the two genes revealed that co-bursting of divergent transcripts is disfavored, suggesting that at higher temporal resolution the oppositely oriented core promoters may compete at the level of transcription initiation. The mechanisms underlining this competition are unclear, but may include steric hindrance and/or local DNA structure alterations, where supercoiling from transcription in one direction impacts transcription dynamics of the other^55,56^.

The competitive relationship of transcriptional bursting was unexpected since at the PRO-seq level no significant reduction in sense transcription was observed upon uasTrx upregulation. We speculate that compensatory mechanisms may buffer against reduction in sense transcription in cases where maintenance of normal gene expression is essential. Finally, although divergent transcription is largely concordant in population-based assays^1,2,8-10^, that concordance might be a reflection of overall promoter strength rather than a direct coordination of sense/antisense core promoters.

CTCF at gene promoters has been invoked to facilitate communication with enhancers^16,57^. Nevertheless, CTCF (previously also known as NeP1) was originally shown to function as a direct transcriptional repressor in reporter gene assays^39,58^, either alone or perhaps by aiding the adjacent binding of a distinct repressor molecule^58^. The CTCF function uncovered here is novel and distinct in that it blocks initiation selectively of uasTrx production at hundreds of genes without significantly impacting sense transcription. Whether CTCF inhibits chromatin binding of PIC components directly by steric hindrance, by recruiting co-repressors, or by facilitating the binding of neighboring repressor molecules remains to be determined. Regardless, our study demonstrates that CTCF can play separate and independent roles in both genome architecture and transcriptional regulation, even at sites with architectural connectivity.

In sum, we uncovered a novel role for CTCF as direct and selective repressor of uasTrx production, independently of its architectural functions, which expands CTCF’s role in the control of the non-coding genome.

## Acknowledgments

We are grateful to Hardison, Raj, Lis, and Blobel laboratories for insightful discussions.

## Funding

This work was supported by NIH grants R01 DK054937 to G.A.B., R24 DK106766 to G.A.B. and R.C.H., T32 HL07439 to C.M.S., R01GM121613 to R.C.H, and U01 DK127405 to G.A.B. and A.R.

## Author contributions

J.L. and G.A.B. conceived the study and designed experiments. J.L. and C.A.K. performed ChIP-seq experiments; J.L., M.W.V., and B.M.G. performed ChIP-seq analysis. J.L., C.M.S, and A.H. performed CRISPR editing experiments and 4C experiments. 4C results were analyzed by J.L. and S.Z. J.M.L. performed PRO-seq experiments; J.L. and Z.Z. analyzed PRO-seq results with advice from J.M.T. and J.T.L. C.M.S., M.G., and A.H. performed single-molecule FISH experiments, with data analyzed by C.M.S., J.L., and A.C. under the supervision of A.R. J.L. and G.A.B. wrote the manuscript with input from all authors.

## Competing interests

The authors declare no competing interests.

## Data and materials availability

All sequencing and processed data have been deposited at GEO under accession GSE173442, GSE173443, GSE173444.

## Methods

### Experiments

#### Cell culture and maintenance

G1E-ER4 is an established murine erythroblast cell line^59^. G1E-ER4 cells were grown in IMDM+15% FBS, penicillin/ streptomycin, Kit ligand, monothioglycerol and erythropoietin in a standard tissue culture incubator at 37ºC with 5% CO_2_. Cells were maintained at density below 1 million/ml at all times. Transient CTCF depletion in G1E-ER4 cells was induced by 1mM auxin in culture. Nascent RNA half-life was assessed by quantifying transcript levels via smFISH and RT-qPCR after transcription blockade for 0h, 4h and 6h with 75uM DRB. HCT-116 cells were cultured in McCoy’s 5A medium supplemented with 10% fetal bovine serum, 2 mM L-glutamine, 100 U ml^−1^ penicillin, and 100 µg ml^−1^ streptomycin at 37°C with 5% CO2.

#### siRNA-mediated CTCF/Nipbl depletion

RNAi was performed in HCT-116 cells as previously described using the same published guide sequences^60^ with a final siRNA concentration of 50 nM (non-targeting control, NIPBL) or 150 nM (CTCF). Cells were harvested after 72 hr treatment.

#### CRISPR-Cas9-mediated genome editing

We performed all CRISPR editing in a previously established Cas9-TagBFP expressing G1E-ER4 cell line to enhance editing efficiency^23^. All sgRNA encoding oligonucleotides were inserted into a retroviral U6-sgRNA-PGK-GFP expression vector^61^ using a BsmBI restriction site and transfected into cells by Amaxa II electroporator (Lonza; program G-016) and Amax II Cell Line Nucleofector Kit (R) (Lonza, VCA-1001). GFP+ cells were sorted by FACS 24h post-transfection, followed by single-cell clone screening and genotyping by Sanger sequencing. All guide RNA sequences were obtained using CRISPR design tool (https://zlab.bio/guide-design-resources)^62^. Guide sequences are listed in Extended Data Table 2.

#### PRO-seq library preparation

PRO-seq libraries in G1E-ER4 was performed as described previously^23^. For each library, 50 million cells were used with 2 million Drosophila Schneider 2 (S2) cells added as spike-in to control for potential global bias associated with library scaling. Fragments longer than 140bp from the PCR-amplified library were selected and sequenced (2×75bp) on the Illumina NextSeq 500 platform according to manufacturer’s instructions to a depth of ∼100 million/library.

PRO-seq libraries in HCT-116 were performed by the Nascent Transcriptomics Core at Harvard Medical School, Boston, MA. Specifically, aliquots of frozen (−80°C) permeabilized cells were thawed on ice and pipetted gently to fully resuspend. For each sample, 1 million permeabilized cells were used, with 50,000 permeabilized Drosophila S2 added for normalization. Nuclear run on assays and library preparation were performed as described^63^ with following modifications: 2X nuclear run-on buffer consisted of 10 mM Tris (pH 8), 10 mM MgCl2, 1 mM DTT, 300mM KCl, 40uM/ea biotin-11-NTPs (Perkin Elmer), 0.8U/uL SuperaseIN (Thermo), 1% sarkosyl. Run-on reactions were performed at 37°C. Adenylated 3’ adapter was prepared using the 5’ DNA adenylation kit (NEB) and ligated using T4 RNA ligase 2, truncated KQ (NEB, per manufacturer’s instructions with 15% PEG-8000 final) and incubated at 16°C overnight. 180uL of betaine blocking buffer (1.42g of betaine brought to 10mL with binding buffer supplemented to 0.6 uM blocking oligo (TCCGACGATCCCACGTTCCCGTGG/3InvdT/)) was mixed with ligations and incubated 5 min at 65°C and 2 min on ice prior to addition of streptavidin beads. After T4 polynucleotide kinase (NEB) treatment, beads were washed once each with high salt, low salt, and blocking oligo wash (0.25X T4 RNA ligase buffer (NEB), 0.3uM blocking oligo) solutions and resuspended in 5’ adapter mix (10 pmol 5’ adapter, 30 pmol blocking oligo, water). 5’ adapter ligation was per Reimer but with 15% PEG-8000 final. Eluted cDNA was amplified 5-cycles (NEBNext Ultra II Q5 master mix (NEB) with Illumina TruSeq PCR primers RP-1 and RPI-X) following the manufacturer’s suggested cycling protocol for library construction. A portion of preCR was serially diluted and for test amplification to determine optimal amplification of final libraries. Pooled libraries were sequenced using the Illumina NovaSeq platform.

#### RNA extraction, cDNA synthesis and RT-qPCR

Cells were harvested in buffer RLT Plus (Qiagen, Cat # 1053393) with lysate homogenized using QIAshredders (Qiagen, Cat # 79656), followed by RNA purification with RNeasy Mini Kit that included an on-column DNase treatment step (Qiagen, Cat. #74106). Complementary DNA (cDNA) was synthesized with iScript Supermix (Bio-Rad, Cat. #1708841). Quantitative polymerase chain reaction (qPCR) was performed using Power SYBR Green kit (Invitrogen; 4368577) with signals detected by ViiA7 System (Life Technologies). Primers used for RT-qPCR are listed in Extended Data Table 3.

#### ChIP-seq library preparation

Chromatin immunoprecipitation (ChIP) was performed as previously described^64^. Antibodies include: CTCF (Millipore; 07-729), POLR2A (Cell Signaling; Cat#14958), IgG from rabbit serum (Sigma; 15006). Quantitative polymerase chain reaction (qPCR) was performed using Power SYBR Green kit (Invitrogen; 4368577) with signals detected by ViiA7 System (Life Technologies). ChIP-seq libraries were prepared using Illumina’s TruSeq ChIP sample preparation kit (Illumina, Cat#IP-202-1012) according to manufacturer’s specifications, with the addition of size selection (left side at 0.9x, right side at 0.6x) using SPRIselect beads (Beckman Coulter, Cat#B23318). Library size was determined (average 351 bp, range 333-372 bp) using the Agilent Bioanalyzer 2100, followed by quantitation using real-time PCR using the KAPA Library Quant Kit for Illumina (KAPA Biosystems; Cat#KK4835). Libraries were then pooled and sequenced (1×75bp) on the Illumina NextSeq 500 platform according to manufacturer’s instructions. Bclfastq2 v 2.15.04 (default parameters) was used to convert reads to fastq. Primers used for RT-qPCR are listed in Extended Data Table 4.

#### 4C-seq sample preparation

The 4C experiments were performed as previously described using DpnII and Csp6I as restriction enzymes^65,66^. Sequencing was done on Illumina Hiseq 2000 genome sequencer with reads mapped onto mm9. Reads mapping to multiple fragment ends were removed, and 4C coverage was computed by averaging mapped reads in running windows of 41 fragment ends. Amplification primers for each view point are listed in Extended Data Table 5. Quality of all libraries meet the previously described standards^66^ based on the *cis*/overall ratio and the percentage of covered fragends within 0.2Mb window around the viewpoints.

#### smFISH imaging

Single-molecule RNA FISH was performed as previously described^45,67^. All sense probes used were complementary to introns of gene of interest and are listed in Extended Data Table 6. Briefly, cells were fixed in 1.85% formaldehyde for 10 min at room temperature, and stored in 70% ethanol at 4ºC. Pools of fluorophore-conjugated FISH probes were hybridized to samples overnight, followed by DAPI staining and washes performed in suspension. Cells were cytospun onto slides for imaging on a Nikon Ti-E inverted fluorescence microscope using a 100x Plan-Apo objective (numerical aperture of 1.43), a cooled CCD camera (Pixis 1024B from Princeton Instruments), and filter sets SP102v1 (Chroma), SP104v2 (Chroma), and 31000v2 (Chroma) for Cy3, Atto647N, and DAPI, respectively. Slides were imaged in 36 optical z sections at intervals of 0.35 microns with 1 s exposure time for Cy3/Atto647N and 35 ms for DAPI.

### Analysis

#### PRO-seq quantification

Read alignment and identification of active transcripts have been described in detail previously^23^. An arbitrary window of +200bp relative to Refseq-annotated TSS to −500 bp relative to TES (transcription end site) was used to quantify sense transcript levels to avoid any confounding effects associated with promoter-proximal pausing. A window of −1000bp to +200bp relative to TSS was selected to quantify uasTrx changes unless noted otherwise. Differential expression analysis was performed using paired DESeq2 method^68^ with FDR<0.05 & fold-change>1.5 as thresholds. Each up-regulated uasTrx in G1E-ER4s was confirmed visually to rule out false positives such as run-throughs from nearby up-regulated genes. For analysis of PRO-seq datasets published in Rao et al. 2017, only active genes identified by the authors were included for characterization.

The start and end sites of uasTrx were annotated as follows: 1) Reads less than 100bp long were extended to 100bp from the 3’ end to “smooth over” PRO-seq signals. 2) Regions overlapping any known transcripts were masked. 3) Global averaged sequencing depth was obtained by dividing all mapped reads over the entire genome. 4) Unbroken regions starting within 500bp of the annotated TSSs on the antisense strand and with sequencing depth exceeding global average were counted as part of uasTrx and taken into consideration for length estimates.

#### RNA-seq quantification

A window of −2000bp to −50bp relative to annotated TSSs was used to quantify uasTrx in unstranded RNA-seq datasets published in Nora et al. 2017 to minimize inclusion of sense signals. DESeq2 was applied to read count matrix to evaluate differential expression between groups.

#### ChIP-seq analysis

Bowtie 1.1.0 was used to align sequences to the mm9 reference genome^69^. Reads with more than one mismatch or multiple alignments were excluded. Significantly enriched regions were called using MACS2 version 2.1.0^70^ with the following parameters: p = 10 5, extsize = 300 and local lambda = 100,000 using whole-cell extract input controls. Reads for the bigwigs were RPM normalized.

#### smFISH image analysis

Nuclear boundaries were segmented manually from DAPI images, with RNA spots localized and quantified using custom software written in MATLAB^71^. Transcription sites were identified by bright nuclear intron spots; fluorescence intensities of transcription sites were determined by 2D Gaussian fitting on processed image data. Subsequent analysis was performed in R. To identify sense and antisense co-transcription status, a wide range of sense-antisense distance thresholds were tested, ranging from 1 pixel (our resolution limit) to 10 pixels (1.3µm). Almost all distance thresholds yielded similar results. Results shown in Fig. 3 and Extended Data Fig. 11 are based on distance threshold of 3 pixels (0.39µm).

#### Gene ontology analysis

Gene ontology (GO) analysis was performed using PANTHER overrepresentation test (release 20210224) against all Mus musculus genes in the database as background. The Fisher’s exact test was performed with FDR correction. GO Ontology database DOI: 10.5281/zenodo.4495804 (released 2021-02-01).

#### Metaplots

All metaplots were generated as previously described^42^ and show estimated average signals and the 87.5 and 12.5 percentiles obtained from bootstrapping.

**Extended Data Fig.1.**
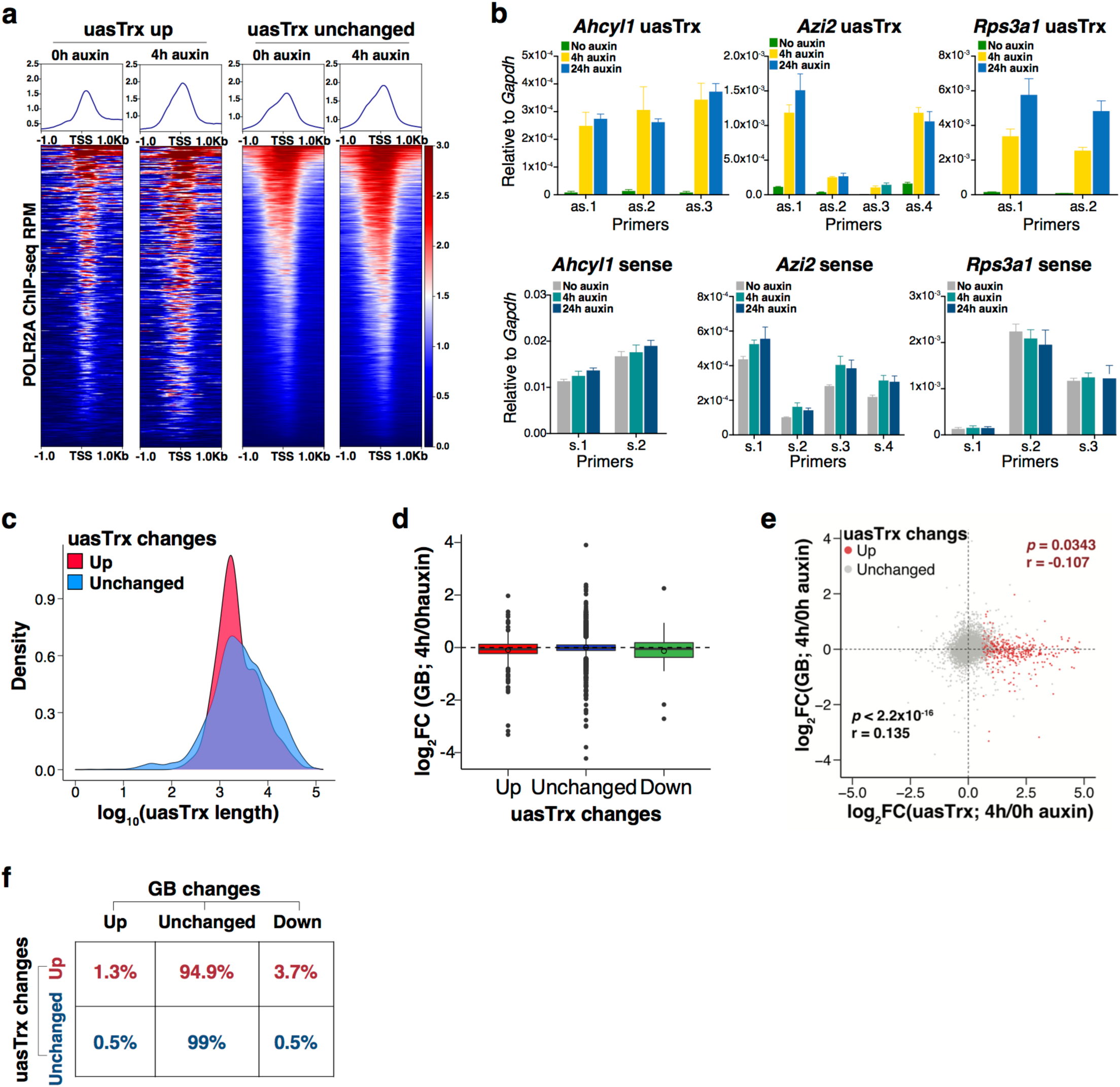
Transient CTCF depletion leads to widespread uasTrx up-regulation at divergent promoters. **a**, Row-linked heatmaps showing POLR2A occupancy at active promoters, grouped by antisense changes (up n= 376; unchanged n=9,632) upon CTCF depletion, sorted by occupancy level, and shown with respect to sense orientation. **b**, Top, RT-qPCR of uasTrx at indicated loci at indicated time points after CTCF depletion. Transcripts were normalized to *Gapdh* (error bar: SEM; n=3-4). Bottom, same as top but quantifying nascent sense transcripts. **c**, Distribution of uasTrx lengths, grouped by changes in response to CTCF depletion. **d**, Log-transformed PRO-seq fold changes in GB after CTCF depletion, grouped by uasTrx changes. **e**, Scatterplot comparing transcriptional changes in GB versus uasTrx. Data points grouped and colored based on uasTrx changes. **f**, Transcriptional changes in uasTrx and GB after CTCF depletion.

**Extended Data Fig.2.**
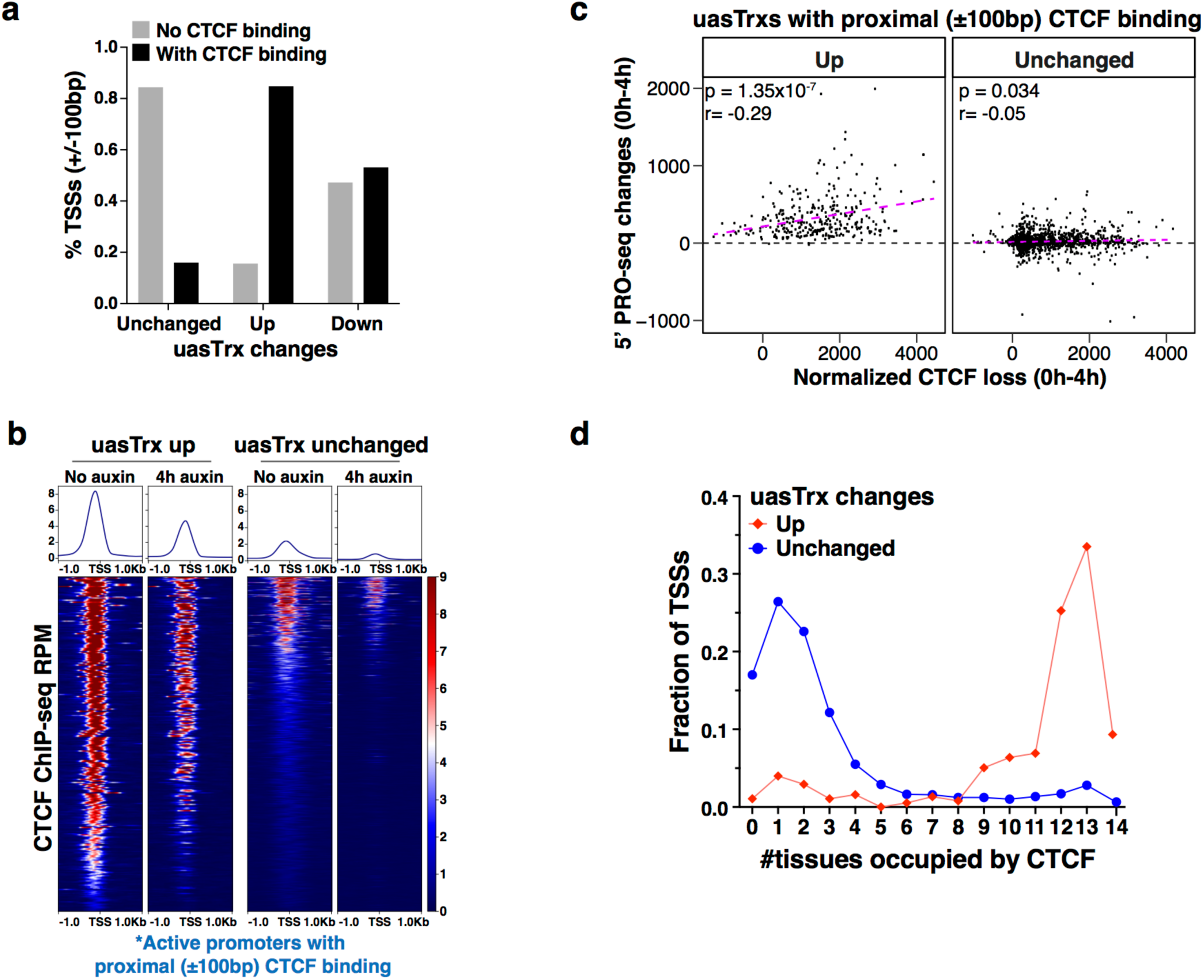
CBSs proximal to activated uasTrx exhibit distinct features. **a**, Percentage of promoters with and without proximal (±100bp) CBSs as a function of uasTrx changes. **b**, Heatmaps showing CTCF occupancy at active promoters with proximal (±100bp) CTCF binding (up n=319; unchanged n=1,527), sorted by occupancy level, and shown with respect to sense orientation. **c**, Correlation between 5’ PRO-seq changes and CTCF loss at uasTrx with proximal (±100bp) CTCF binding. Linear regression line shown in magenta. *P* value was calculated by Spearman rank correlation test; r is the correlation coefficient. **d**, Fraction of TSSs detected in the indicated numbers of mouse tissues where CTCF binds in proximity (within ±100bp), grouped by uasTrx changes.

**Extended Data Fig.3.**
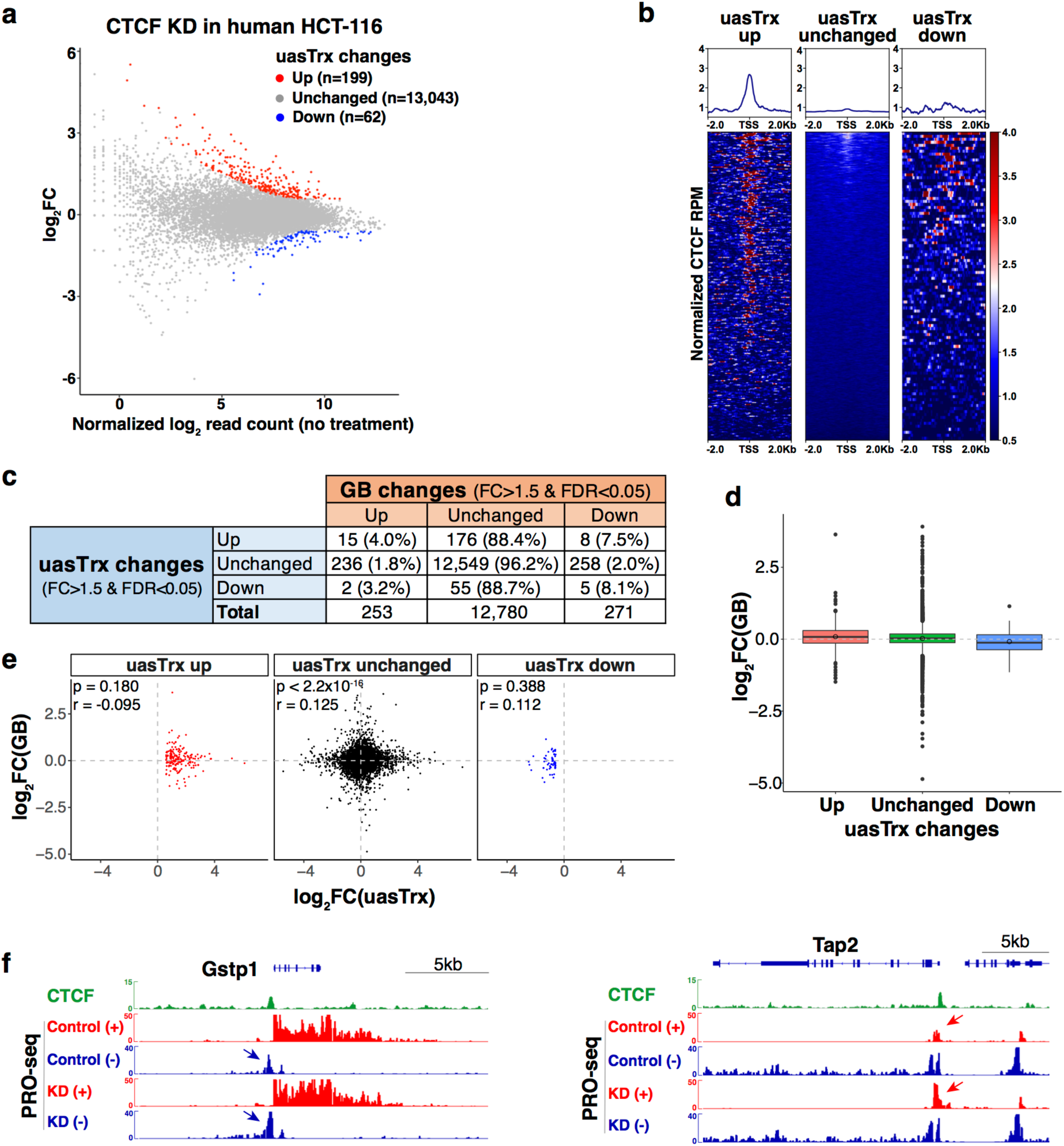
Transient CTCF depletion in human HCT-116 leads to similar antisense transcriptional changes. **a**, PRO-seq MA plot of control versus CTCF-depleted cells on the antisense strand (−1000bp to +200 relative to annotated TSS) in human HCT-116 cells. Differentially expressed transcripts highlighted in color. **b**, Row-linked heatmaps showing CTCF occupancy at active promoters, grouped by uasTrx changes, sorted by binding enrichment levels, and shown with respect to sense orientation. **c**, Transcriptional changes in uasTrx and GB after CTCF depletion. **d**, Boxplot showing log-transformed PRO-seq fold changes in GB. **e**, Scatterplot showing log-transformed PRO-seq fold changes in GB and uasTrx. **f**, Brower views of CTCF ChIP-seq (mm9 liftover from Rao et al., 2014) and PRO-seq signals at *Gstp1* and *Tap2* loci. Arrows highlight location of CTCF-repressed uasTrx. Arrow color indicates uasTrx strandedness. KD, knockdown.

**Extended Data Fig.4.**
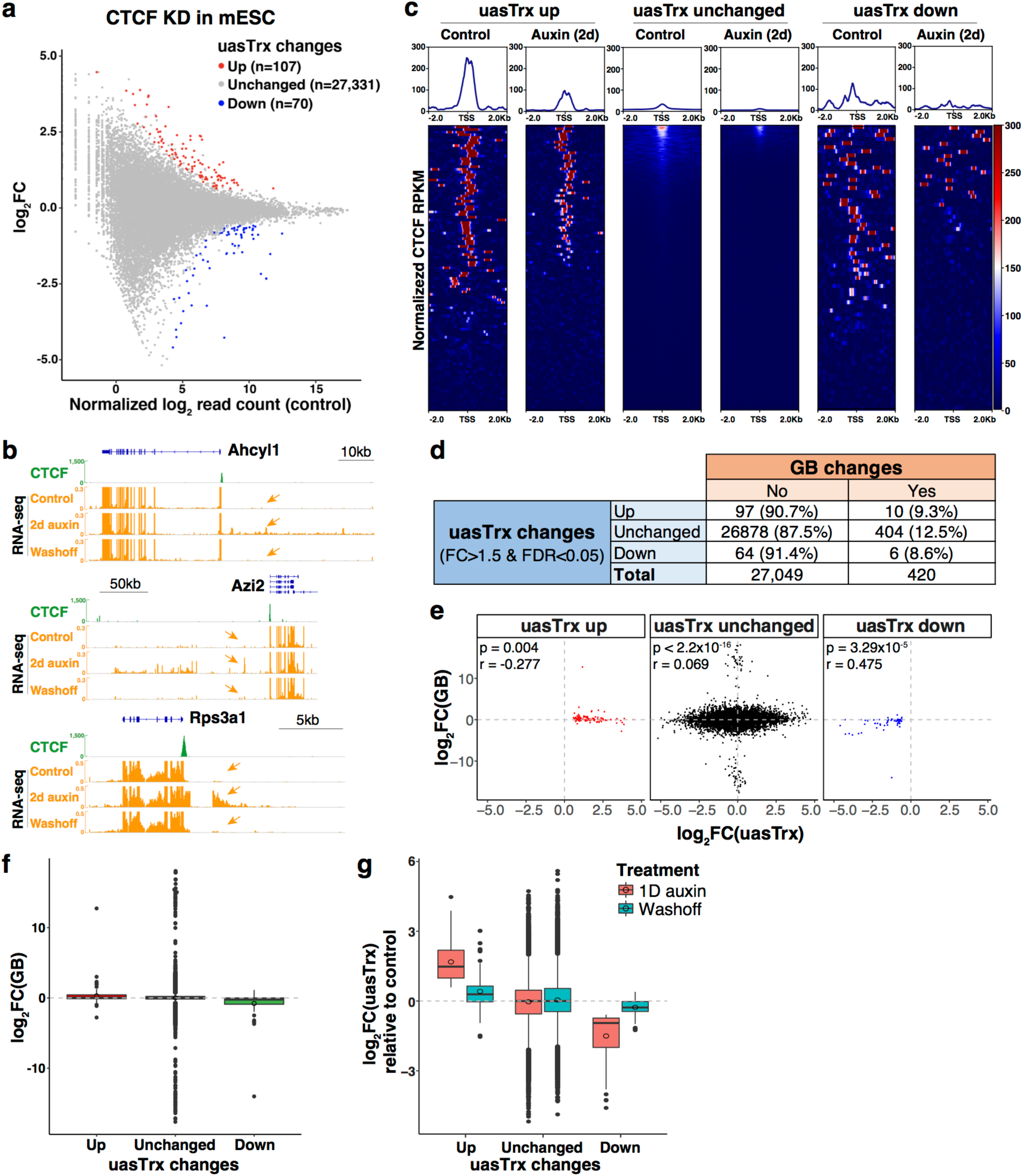
Transient CTCF depletion in mESC leads to similar antisense transcriptional changes. **a**, RNA-seq MA plot of control versus CTCF-depleted cells on the antisense strand (−1000bp to +200 relative to annotated TSS) in mESC. Differentially expressed transcripts highlighted in color. **b**, Brower views of CTCF ChIP-seq and RNA-seq signals at *Ahcyl1, Azi2* and *Rps3a1* loci. Arrows highlight signals upstream of TSS indicative of antisense transcription. **c**, Row-linked heatmap showing CTCF occupancy at active promoters, grouped by uasTrx changes and shown with respect to **s**ense orientation. **d**, Transcriptional changes in uasTrx and GB after CTCF depletion. **e**, Correlation between uasTrx and GBs changes in RNA-seq upon CTCF depletion. *P* value was calculated by Spearman rank correlation test; r is the correlation coefficient. **f**, Log-transformed RNA-seq fold changes in GB after CTCF depletion over control. **g**, Log-transformed RNA-seq fold change in uasTrx in indicated conditions over control. Note the repression of elevated uasTrx after auxin washoff.

**Extended Data Fig.5.**
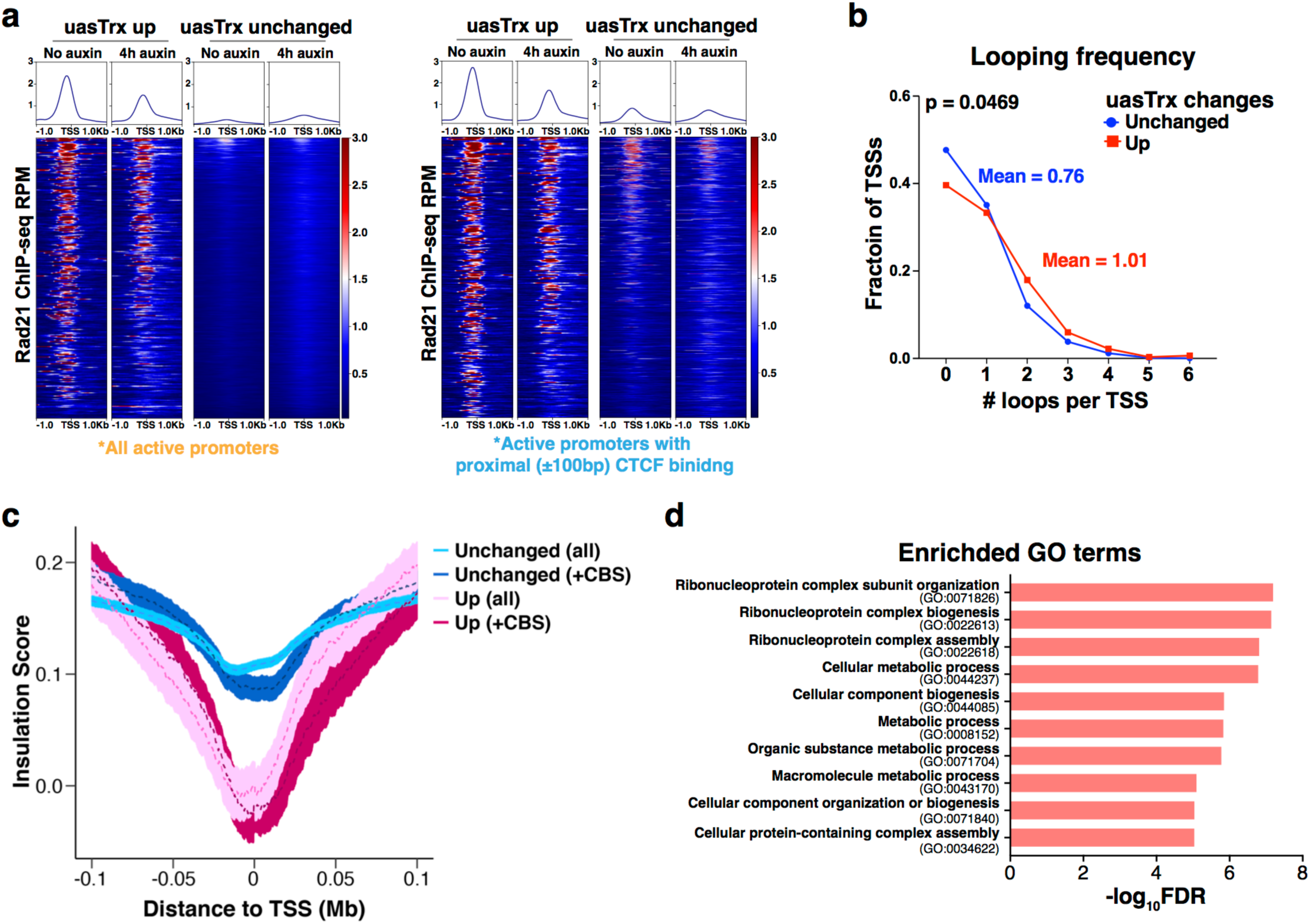
Affected promoters are associated with architectural features. **a**, Left, row-linked heatmaps showing Rad21 occupancy at all activated (n=376) and unaffected (n=9,632) active promoters, grouped by CTCF depletion-elicited uasTrx changes, sorted in the same order as **Fig. 1e**, and shown with respect to sense orientation. Right, same as left except only plotting those with proximal (±100bp) CTCF binding (up n=319; unchanged n=1,527). **b**, Distribution of looping frequencies of up-regulated versus unchanged uasTrx with proximal (±100bp) CTCF binding. *P* value calculated by Wilcoxon signed-rank test. **c**, Averaged insulation score centered at annotated TSS over 0.2Mb window, plotted with respect to sense orientation, and grouped by uasTrx changes and whether CTCF binds proximally (±100bp; “+CBS”). **d**, Gene ontology terms enriched at genes with activated uasTrx.

**Extended Data Fig.6.**
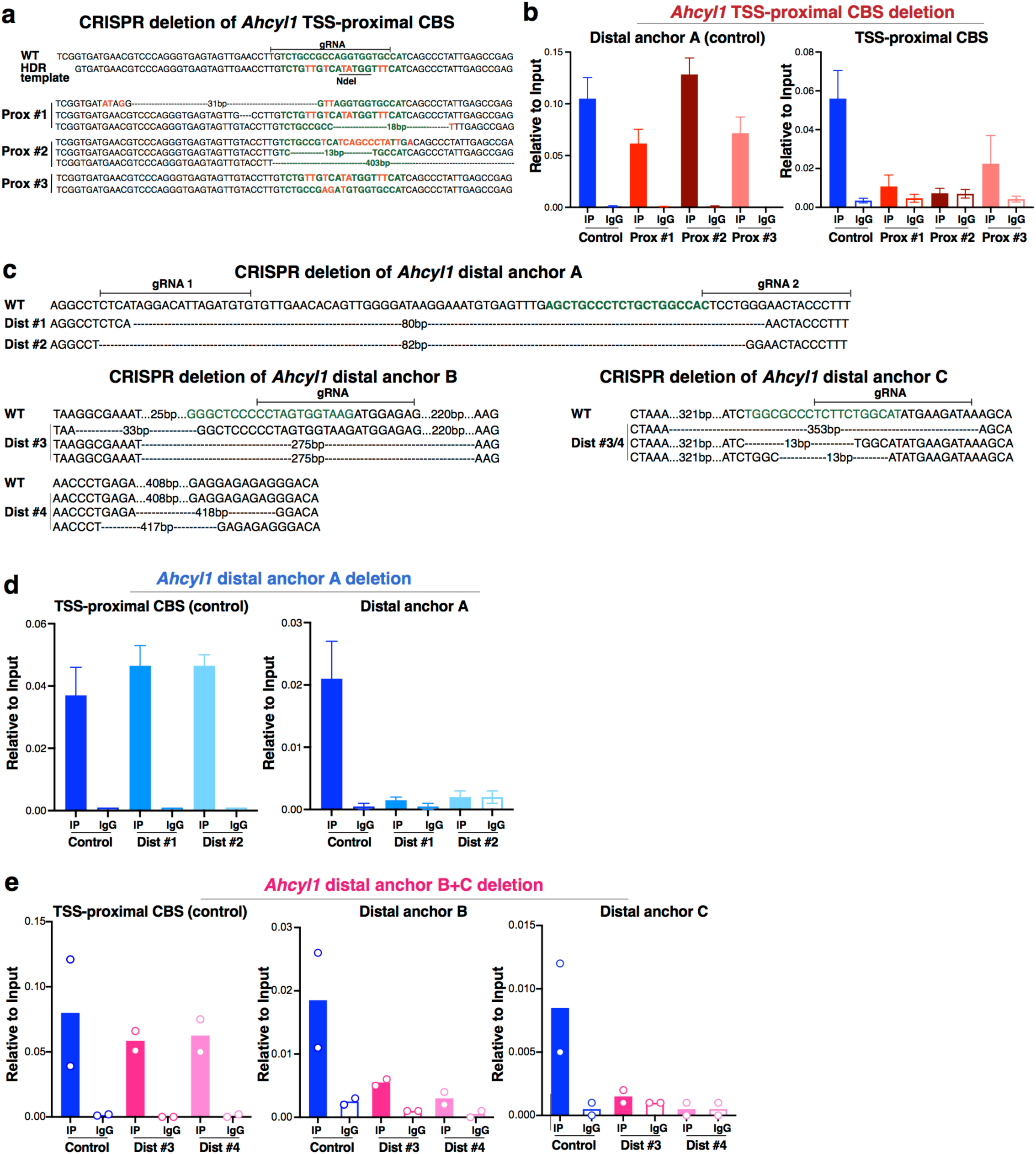
CRISPR/Cas9-mediated genome editing disrupts CTCF binding at *Ahcyl1*. **a**, Genotype of edited clones shown in **Fig. 2c**. Predicted CTCF motif highlighted in green. **b**, Left, CTCF ChIP-qPCR showing abrogation of CTCF binding at TSS-proximal CBS in mutants shown in **Fig. 2c**. Right, distal CBS served as a control for ChIP efficiency (error bar: SEM; n=3). **c**, Genotype of distally edited clones shown in **Fig. 2c**. Predicted CTCF motif highlighted in green. **d**, Left, TSS-proximal CBS served as a control for ChIP efficiency. Right, CTCF ChIP-qPCR showing abrogation of CTCF binding at distal anchor A in distal clones #1 and 2 shown in **Fig. 2c** (error bar: SEM; n=3). **e**, Left, TSS-proximal CBS served as a control for ChIP efficiency. Middle, CTCF ChIP-qPCR showing abrogation of CTCF binding at distal anchor B in distal clones #3 and 4 shown in **Fig. 2c** (n=2). Right, same as middle but measuring binding at distal anchor C.

**Extended Data Fig.7.**
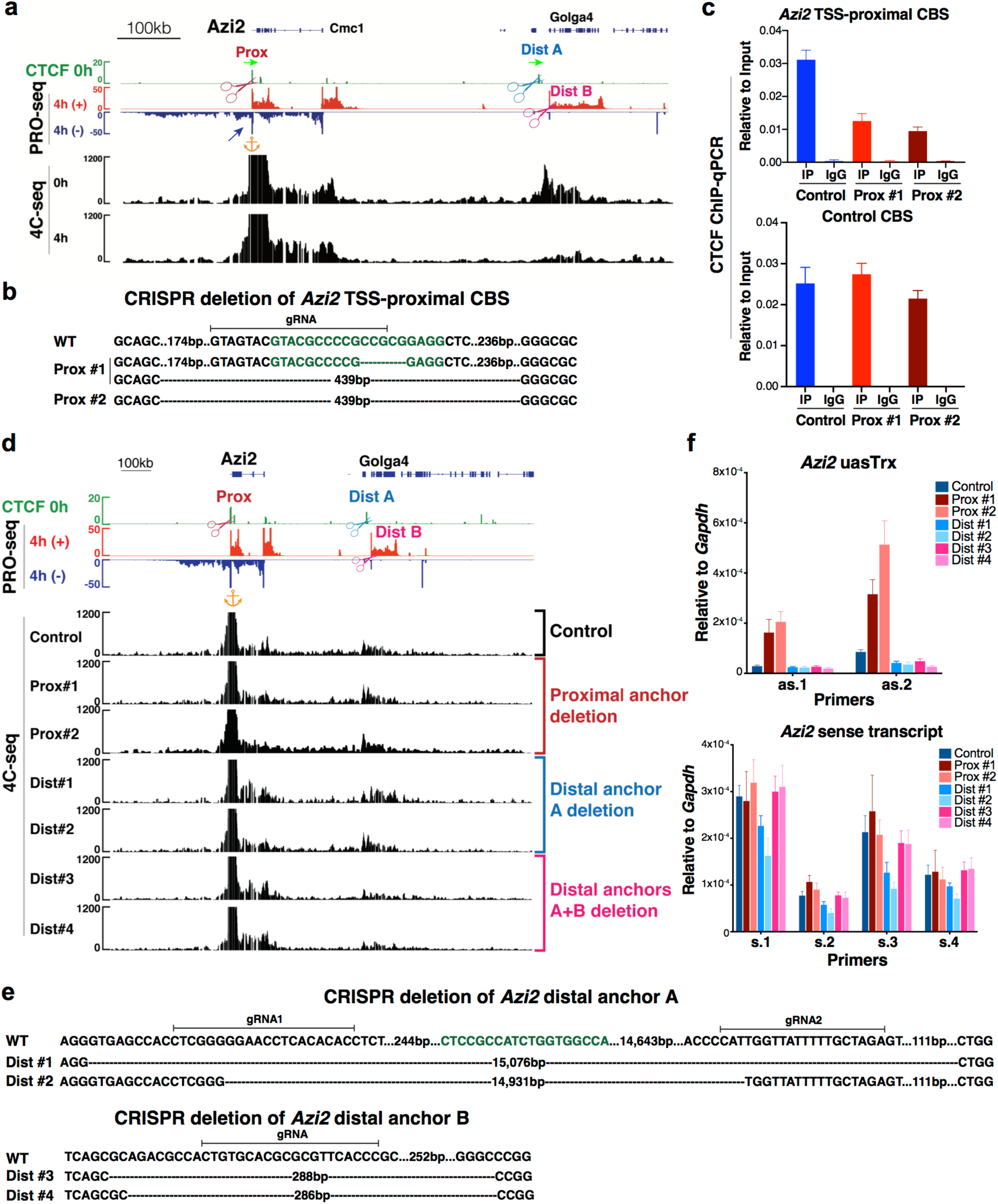
CRISPR/Cas9-mediated deletion of TSS-proximal CBS, but not distal loop anchors, at *Azi2* leads to uasTrx activation. **a**, Genome browser tracks of CTCF ChIP-seq, PRO-seq and 4C-seq at *Azi2* locus, with uasTrx highlighted by dark blue arrow and CRISPR targeted regions indicated by scissors. Green arrows indicate CTCF motif directionality. Orange anchor shows 4C-seq viewpoint. **b**, Genotype of TSS-proximally edited clones. Predicted CTCF motif highlighted in green. **c**, Top, ChIP-qPCR confirming disruption of CTCF binding in mutants (error bar: SEM; n=3). Bottom, ChIP-qPCR at an independent locus controlling for ChIP efficiency (error bar: SEM; n=3). **d**, Left, representative 4C-seq profiles of control/mutant clones with edited regions indicated. Genome browser tracks of bulk CTCF ChIP-seq and PRO-seq shown on top. Similar observations were made in 2-3 independent 4C-seq experiments. Orange anchor indicates 4C-seq viewpoint. Scissor indicates edited region. Middle and right, RT-qPCR of nascent antisense and sense transcripts in WT/mutant clones (error bar: SEM; n=4). **e**, Genotype of mutants with distal anchor(s) disrupted. Predicted CTCF motif highlighted in green. **f**, RT-qPCR of *Azi2* uasTrx and sense primary transcripts in control and edited clones. Transcripts were normalized to *Gapdh* (error bar: SEM; n=3-4). Prox, TSS-proximal CBS. Dist, distal anchor.

**Extended Data Fig.8.**
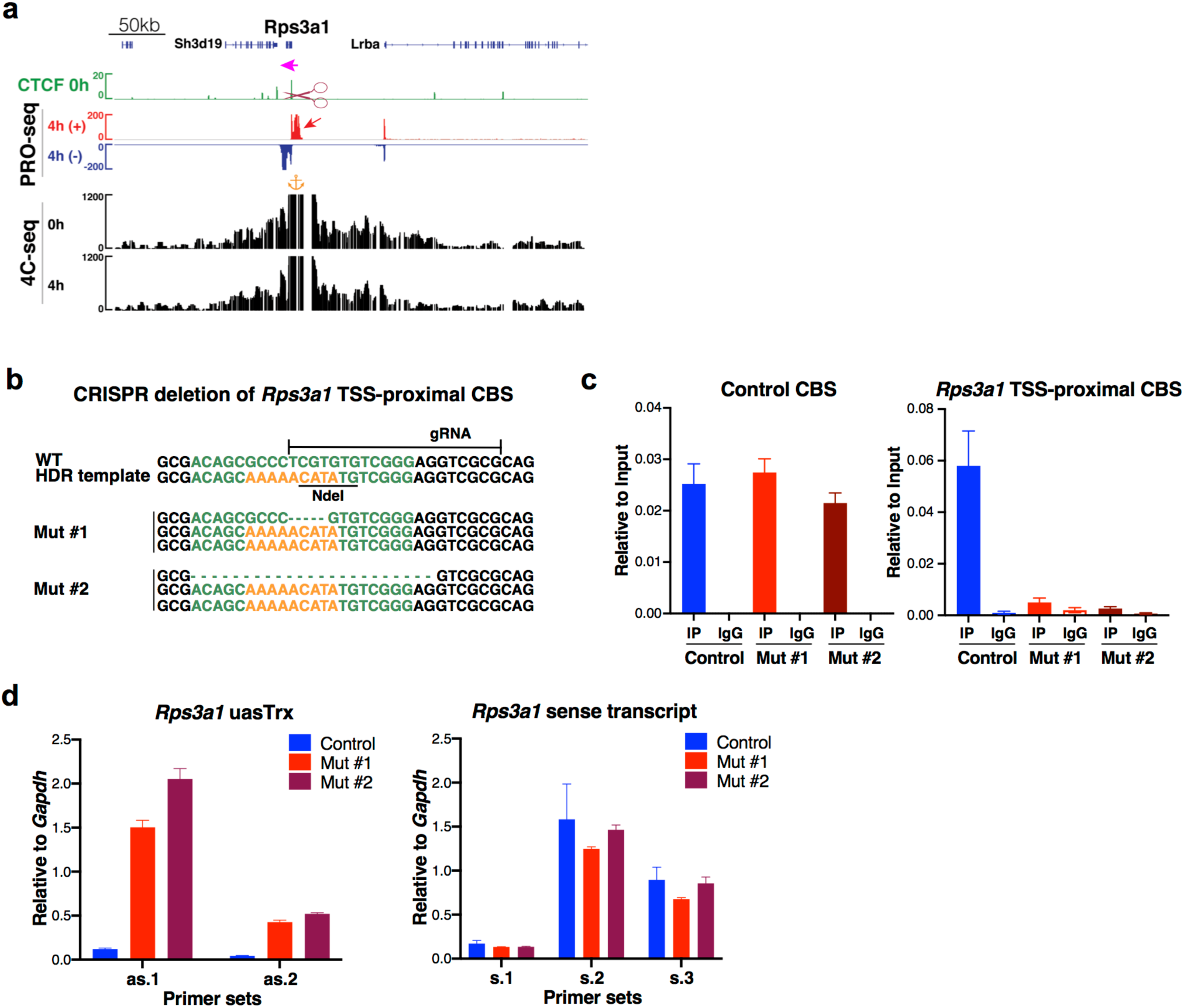
CRISPR/Cas9-mediated deletion of TSS-proximal CBS at *Rps3a1* leads to uasTrx up-regulation. **a**, Genome browser tracks of CTCF ChIP-seq, PRO-seq and 4C-seq at *Rps3a1* locus, with elevated uasTrx highlighted by red arrow and edited region indicated by scissor. Orange anchor indicates 4C-seq viewpoint. Magenta arrow above ChIP-seq track indicates CTCF motif directionality. **b**, Genotype of mutants after CRISPR/Cas9-mediated deletion of TSS-proximal CBS. Predicted CTCF motif highlighted in green. **c**, Left, ChIP-qPCR confirming disruption of CTCF binding in mutant clones (error bar: SEM; n=3). Right, ChIP-qPCR at an independent locus controlling for ChIP efficiency (error bar: SEM; n=3). **d**, RT-qPCR of nascent uasTrx and sense transcripts in control/mutant clones (error bar: SEM; n=3).

**Extended Data Fig.9.**
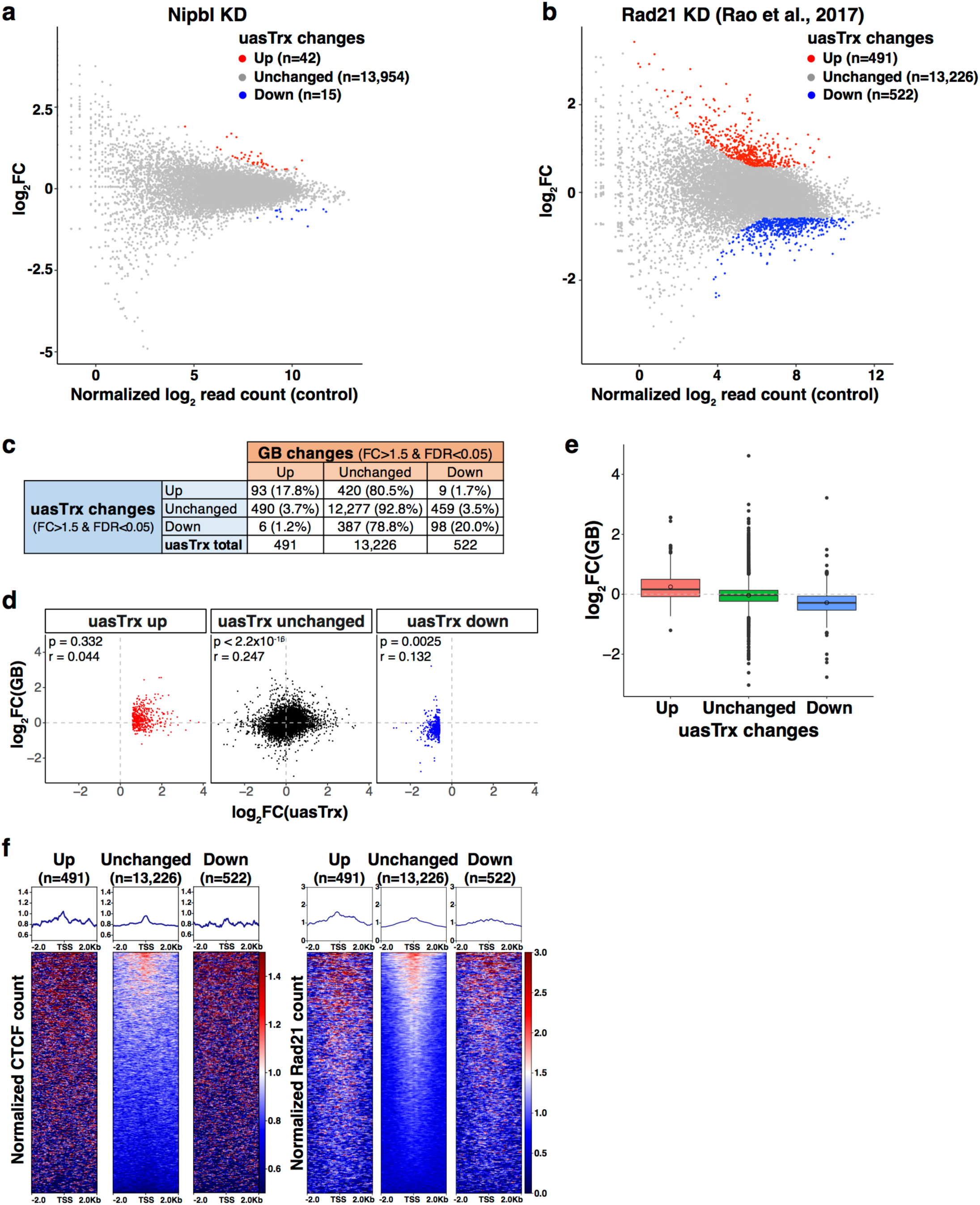
Removal of chromatin-bound cohesin does not recapitulate CTCF-induced uasTrx changes. **a**, PRO-seq MA plot of control versus Nipbl-depleted cells on uasTrx expression (−1000bp to +200 relative to annotated TSS). Differentially expressed transcripts highlighted in color. **b**, Same as **a** but of Rad21-depleted cells. **c**, Table showing the number and percentage of uasTrx and GB changes after Rad21 depletion. **d**, Scatterplot comparing log-transformed 5’ PRO-seq fold changes in uasTrx and GB. *P* value was calculated by Spearman rank correlation test; r is the correlation coefficient. **e**, Boxplot showing log-transformed PRO-seq fold changes in GBs after Rad21 depletion. **f**, Left, row-linked heatmap showing CTCF occupancy at active promoters, grouped by uasTrx changes after Rad21 depletion, sorted by occupancy levels, and shown with respect to sense orientation. Right, same as left, but plotting Rad21 occupancy. Note that neither CTCF nor Rad21 is enriched at up-regulated uasTrx.

**Extended Data Fig.10.**
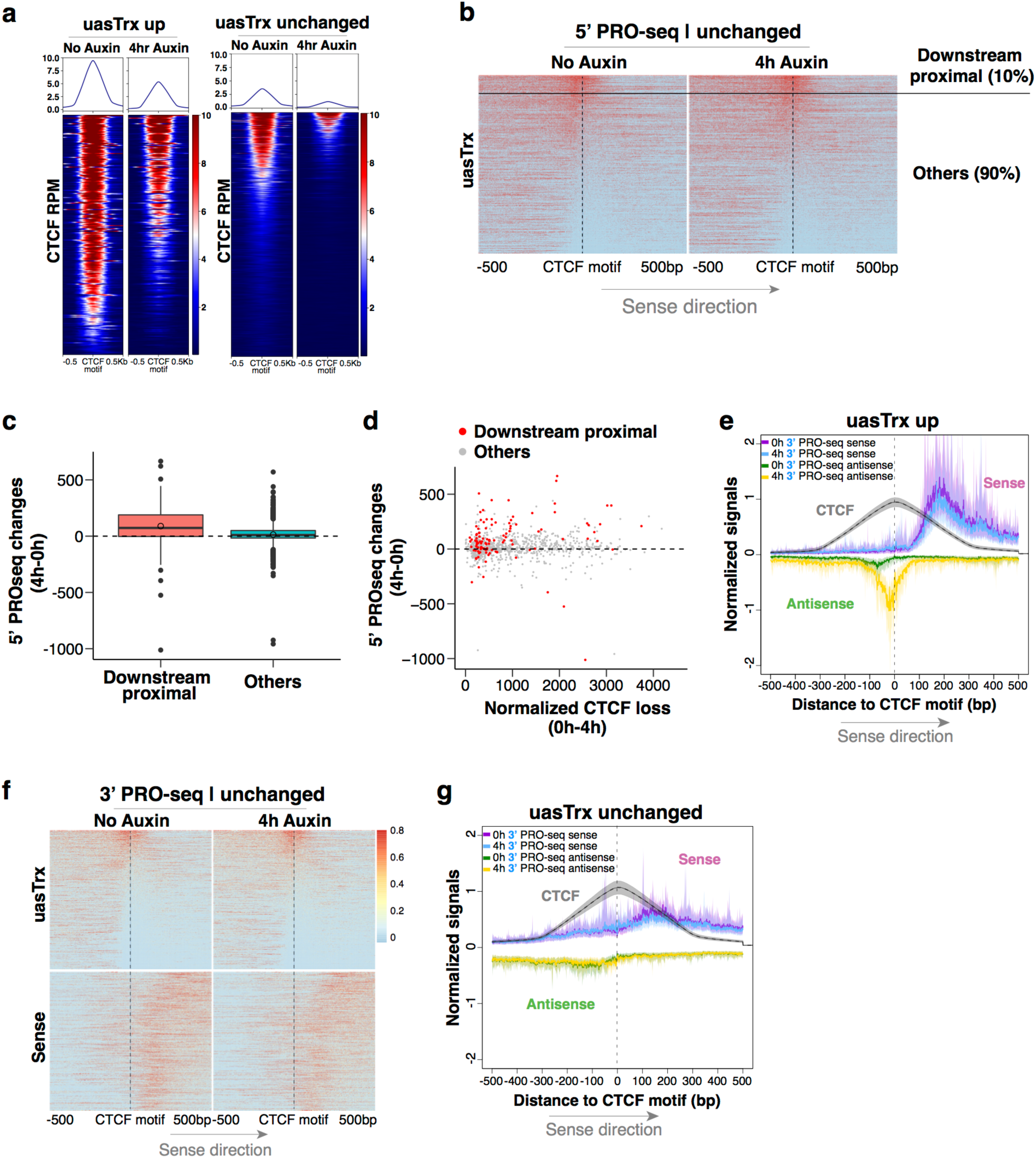
CTCF inhibits antisense transcription initiation through precise positioning. **a**, Left, row-linked CTCF heatmap at affected active promoters that harbor proximal (±100bp) CTCF binding and high-confidence CTCF motif scores (>75), centered at CTCF motifs, grouped by mean signal densities over center 200bp, and shown with respect to sense orientation. Right, same as left but at unaffected active promoters meeting the same CTCF criteria. **b**, 5’ PRO-seq heatmap at unchanged promoters shown in **Fig. 3d** with a portion of sites (10%; “downstream proximal”) manually picked from the rest (“others”), which demonstrate similar CTCF distribution relative to 5’ PRO-seq signals as **Fig. 3b. c**, Related to **b**, plotting PRO-seq changes in uasTrx at unaffected promoters, grouped based on CTCF positioning relative to 5’ PRO-seq signals. **d**, Related to **b**, comparing uasTrx changes and CTCF binding loss at unaffected promoters, grouped based on CTCF positioning relative to 5’ PRO-seq signals. **e**, Metaplot summarizing 3’ PRO-seq and CTCF signals shown in **Fig. 3f**. Solid lines and shades show bootstrapped estimates of average signals and the 12.5/87.5 percentiles, respectively. **f**, Same as **Fig. 3f**, but at unaffected promoters. **g**, Same as **e**, but summarizing **f**.

**Extended Data Fig.11.**
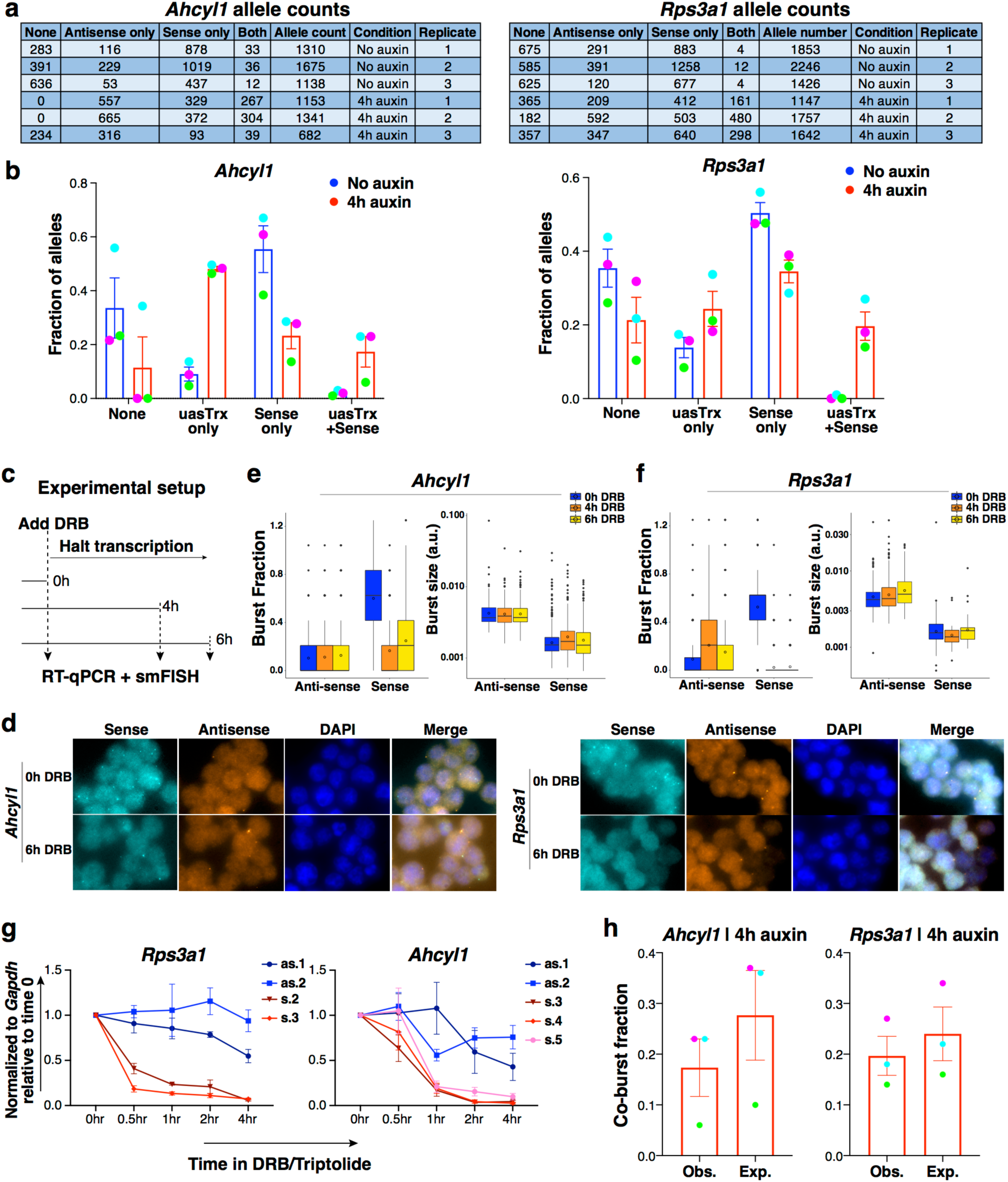
CTCF inhibits antisense burst fraction; sense/antisense co-bursting is disfavored. **a**, Table showing raw smFISH allele counts. **b**, Observed fractions of alleles where sense/antisense burst independently or simultaneously at *Ahcyl1* and *Rps3a1* (error bar: SEM; n=3). Biological replicates matched by dot colors. (**c**, Experimental outline for RNA half-life estimation. **d**, Representative smFISH images before and after DRB treatment at *Ahcyl1* and *Rps3a1*. **e**, Left, box plot showing antisense and sense burst fractions at *Ahcyl1* before and after DRB treatment. Right, same as left but quantifying burst sizes. n=3 biological replicates. *P* values were calculated by two-sample *t*-test. **f**, Same as (**e** but at *Rps3a1*. **g**, RT-qPCR measuring nascent sense and antisense transcript levels at *Ahcyl1* and *Rps3a1* before and after DRB treatment. Transcripts were normalized to *Gapdh* and plotted relative to time 0 (error bar: SEM; n=4). **h**, Predicted and observed co-burst fractions at *Ahcyl1* and *Rps3a1* after 4h auxin treatment (error bar: SEM; n=3). Biological replicates matched by dot colors.

**Extended Data Table 1.** Top gene ontology terms.

| **GO biological process** | **#**<br>**Reference** | **#**<br>**Dataset** | **Expected** | **Fold**<br>**Enrichment** | **+/-** | **raw *P* value** | **FDR** |
| --- | --- | --- | --- | --- | --- | --- | --- |
| Developmental process | 5554 | 54 | 26.02 | 2.08 | + | 4.36E-09 | 6.86E-05 |
| Negative regulation of biological process | 5264 | 46 | 24.66 | 1.87 | + | 4.34E-06 | 1.37E-02 |
| Regulation of multicellular organismal process | 2787 | 31 | 13.06 | 2.37 | + | 3.55E-06 | 1.40E-02 |
| Multicellular organism development | 4788 | 44 | 22.43 | 1.96 | + | 1.96E-06 | 1.54E-02 |
| Anatomical structure development | 5166 | 46 | 24.2 | 1.9 | + | 3.37E-06 | 1.77E-02 |
| System development | 4176 | 39 | 19.56 | 1.99 | + | 7.42E-06 | 1.95E-02 |
| Regulation of developmental process | 2521 | 28 | 11.81 | 2.37 | + | 1.35E-05 | 2.36E-02 |
| Cell differentiation | 3560 | 35 | 16.68 | 2.1 | + | 1.06E-05 | 2.38E-02 |
| Cellular developmental process | 3611 | 35 | 16.92 | 2.07 | + | 1.31E-05 | 2.58E-02 |
| Positive regulation of gene expression, epigenetic | 29 | 4 | 0.14 | 29.44 | + | 1.64E-05 | 2.59E-02 |

**Extended Data Table 2.**
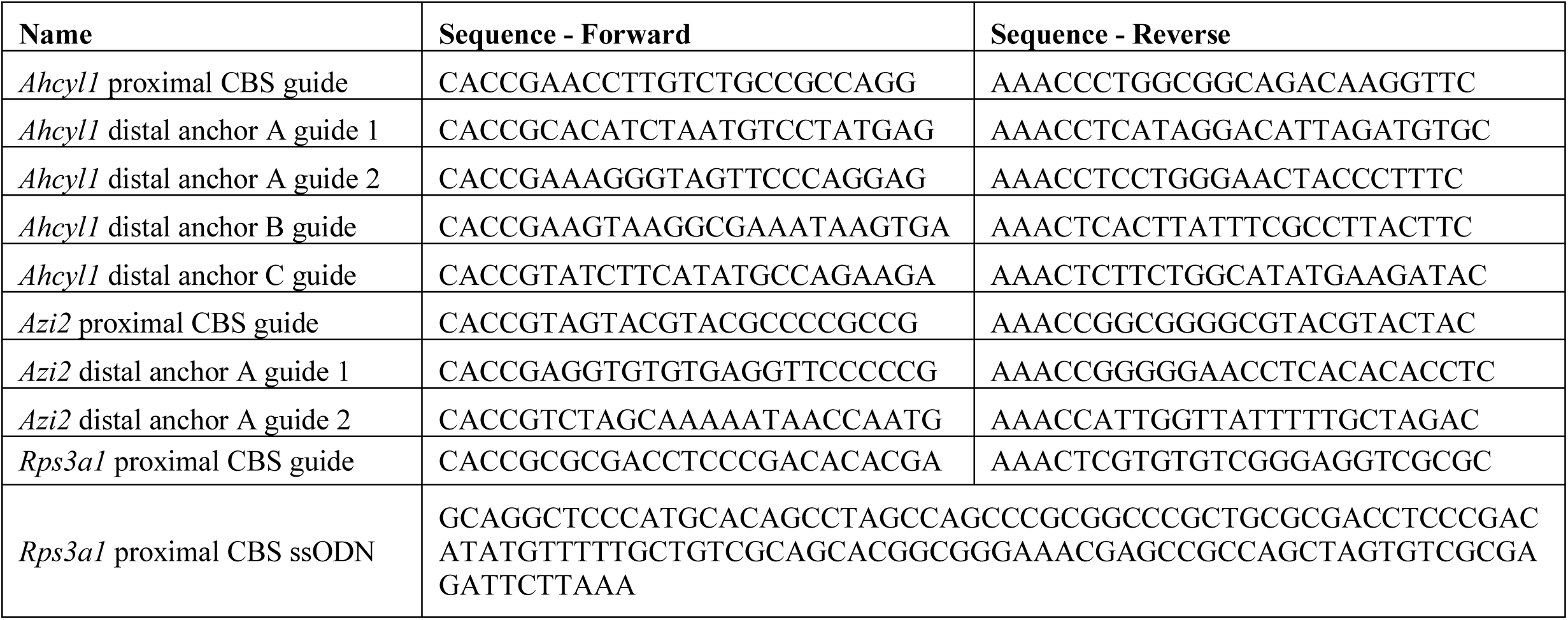
Oligos for sgRNAs.

**Extended Data Table 3.**
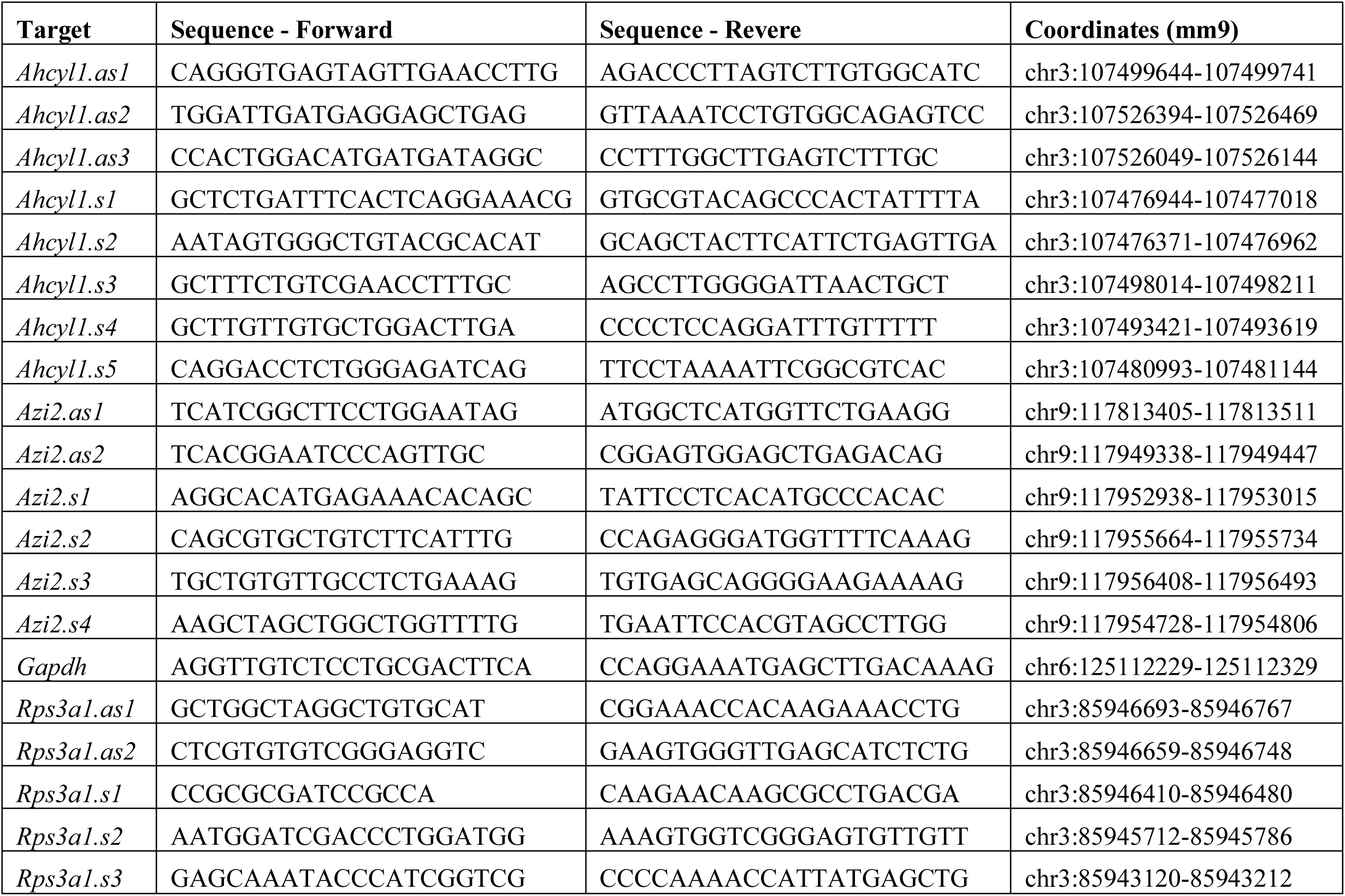
RT-qPCR primers.

**Extended Data Table 4.**
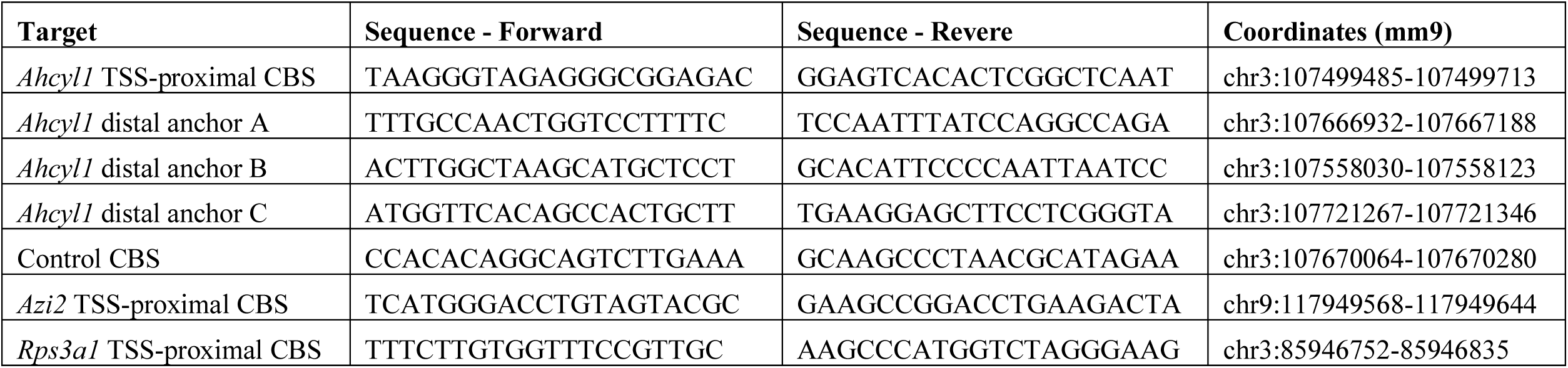
ChIP-qPCR primers.

**Extended Data Table 5.**
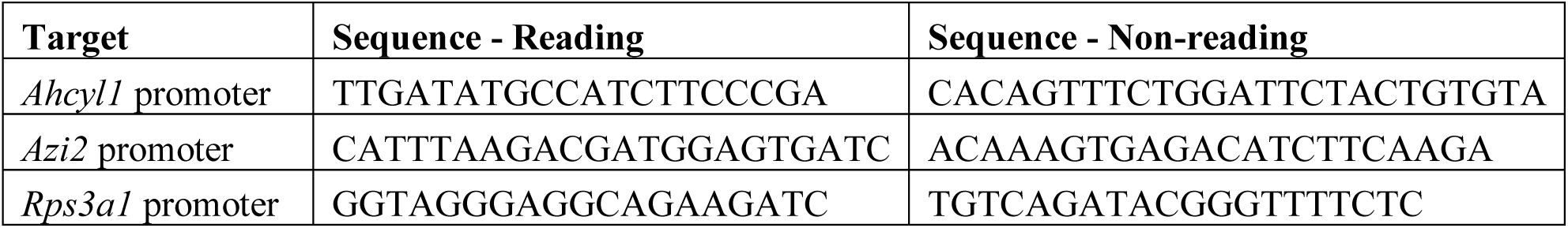
4C primers.

**Extended Data Table 6.**
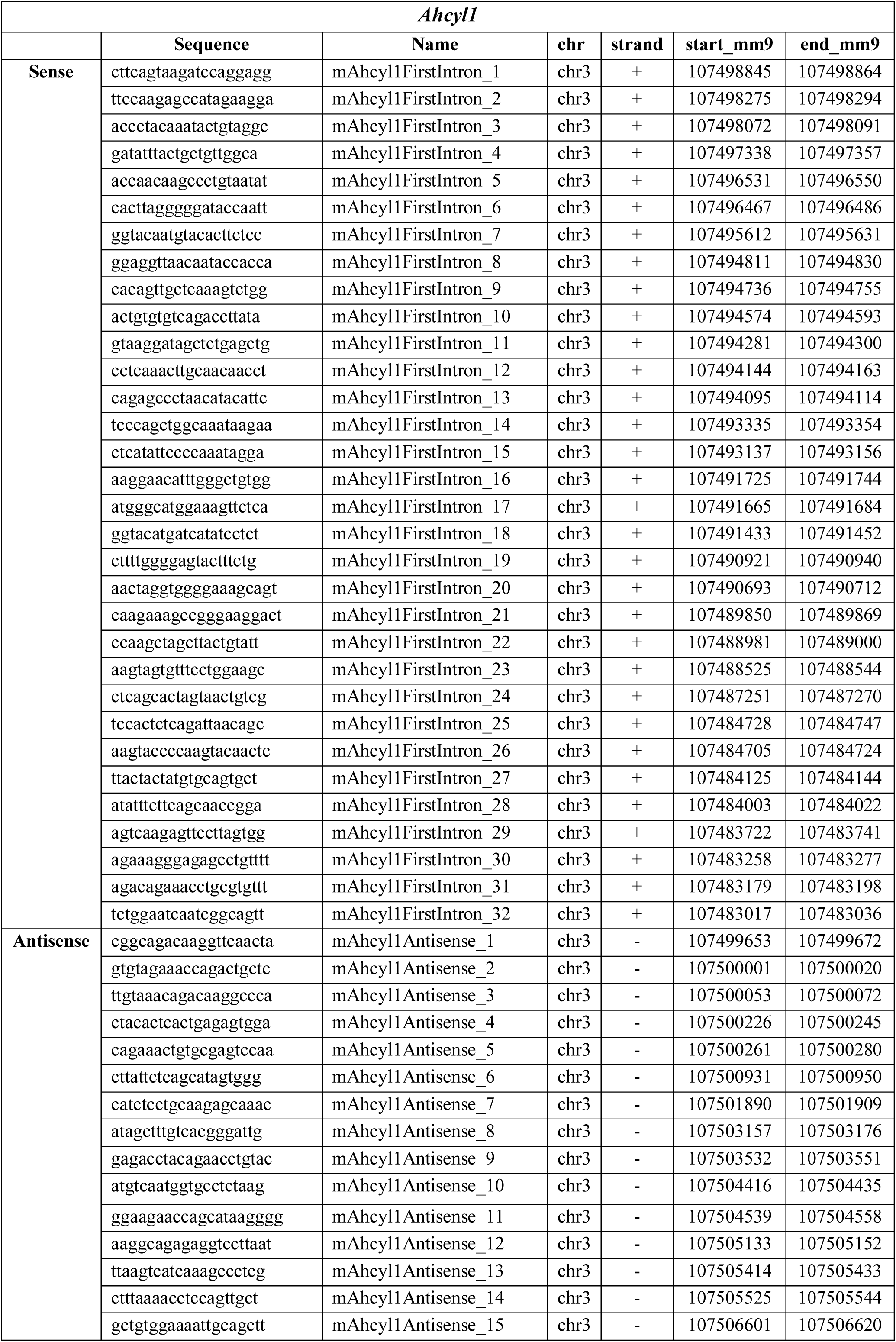
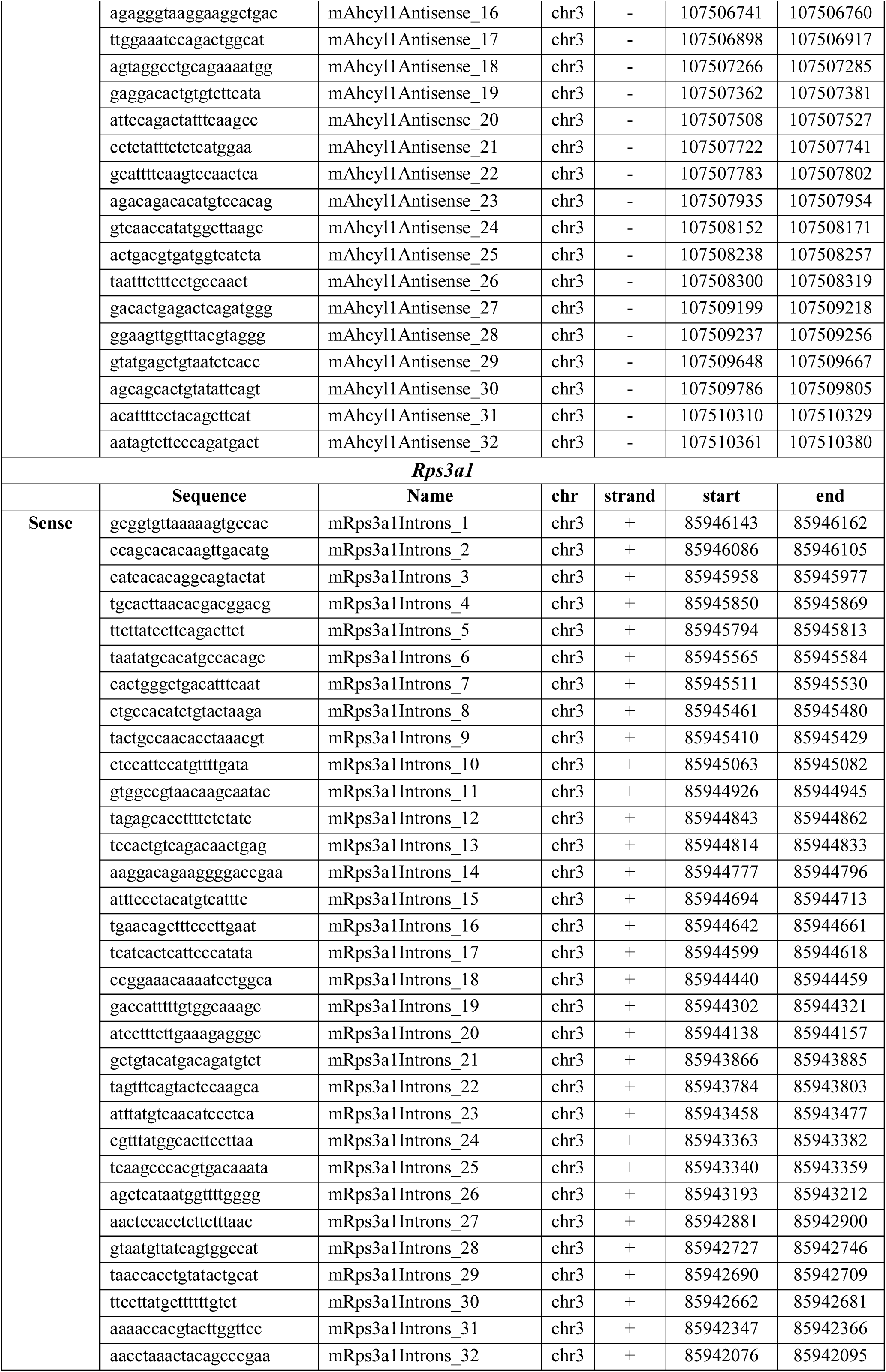
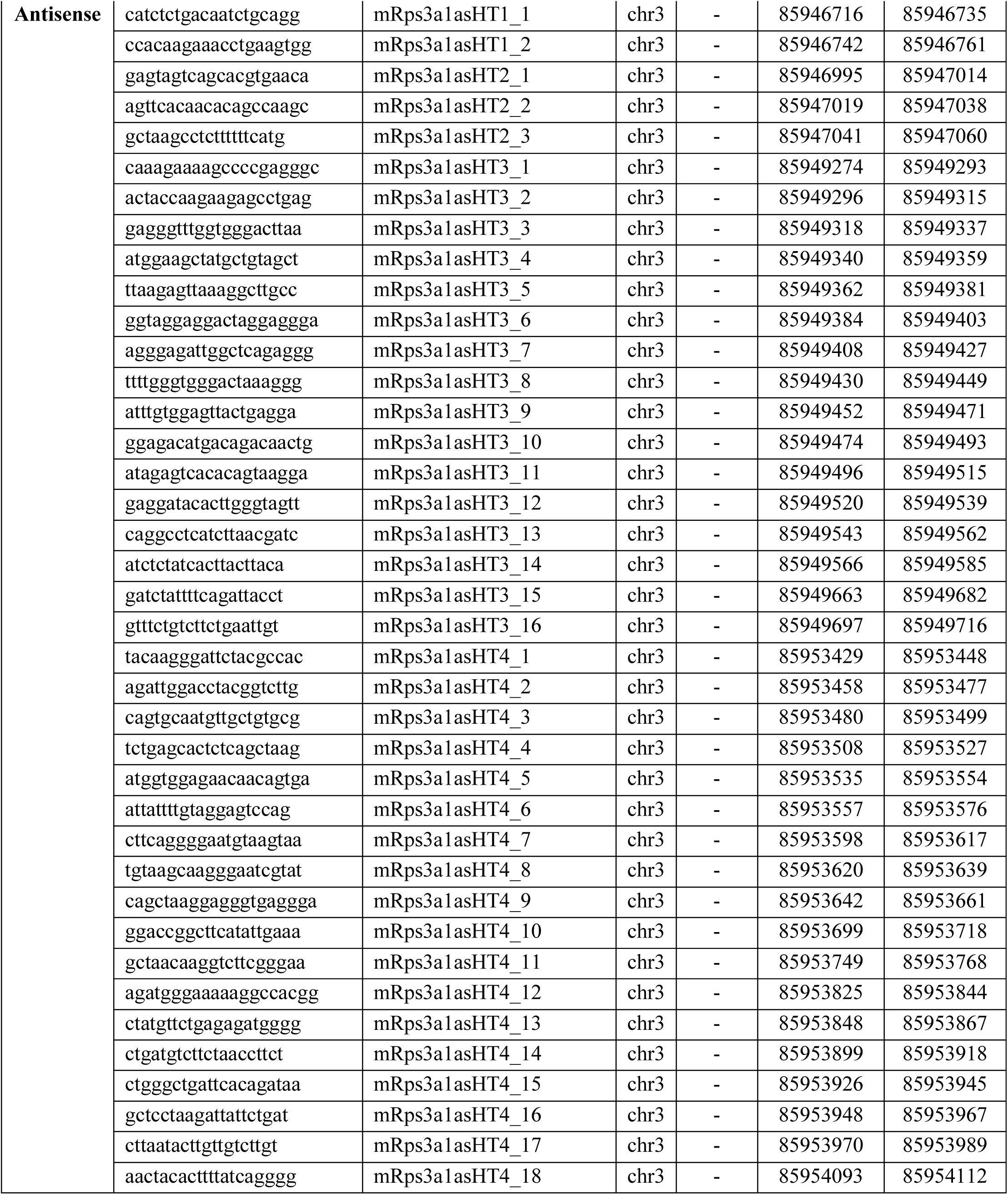
smFISH primers.

## Notes

### Competing Interest Statement

The authors have declared no competing interest.

